# Reassessing face topography in primary somatosensory cortex and remapping following hand loss

**DOI:** 10.1101/2021.07.05.451126

**Authors:** Victoria Root, Dollyane Muret, Maite Arribas, Elena Amoruso, John Thornton, Aurelie Tarall-Jozwiak, Irene Tracey, Tamar R. Makin

**Affiliations:** WIN Centre, University of Oxford, Oxford OX3 9DU, UK; Institute of Cognitive Neuroscience, University College London, London WC1N 3AZ, UK; Department of Psychosis Studies, Institute of Psychiatry, Psychology & Neuroscience, King’s College London, London, UK; Wellcome Trust Centre for Neuroimaging, University College London, London WC1N 3AR, UK; Queen Mary’s Hospital, London, UK

**Author notes:** Correspondence to: Dollyane Muret; Institute of Cognitive Neuroscience, University College London, London WC1N 3AZ, UK. These authors contributed equally to this work (Victoria Root, Dollyane Muret).

**Keywords:** remapping, primary somatosensory cortex, fMRI, face somatotopy, phantom limb pain

## Abstract

Cortical remapping after hand loss in the primary somatosensory cortex (S1) is thought to be predominantly dictated by cortical proximity, with adjacent body parts remapping into the deprived area. Traditionally, this remapping has been characterised by changes in the lip representation, which is assumed to be the immediate neighbour of the hand based on electrophysiological research in non-human primates. However, the orientation of facial somatotopy in humans is debated, with contrasting work reporting both an inverted and upright topography. We aimed to fill this gap in the S1 homunculus by investigating the topographic organisation of the face. Using both univariate and multivariate approaches we examined the extent of face-to-hand remapping in individuals with a congenital and acquired missing hand (hereafter one-handers and amputees, respectively), relative to two-handed controls. Participants were asked to move different facial parts (forehead, nose, lips, tongue) during fMRI scanning. We first report evidence for an upright facial organisation in all three groups, with the upper face and not the lips bordering the hand area. We further found little evidence for remapping of all tested facial parts in amputees, with no significant relationship to the chronicity of their PLP. In contrast, we found converging evidence for a complex pattern of face remapping in congenital one-handers across all facial parts, where the location of the cortical neighbour – the forehead – is shown to shift away from the deprived hand area, which is subsequently activated by the lips and the tongue. Together, our findings demonstrate that the face representation in humans is highly plastic, but that this plasticity is restricted by the developmental stage of input deprivation, rather than cortical proximity.

## Introduction

Our brains capacity to adapt, known as cortical plasticity, is integral to our successful functioning in daily life, as well as rehabilitation from injury. A key model for exploring the extent, and consequences of, cortical plasticity is upper-limb loss (via amputation or congenital absence). Here, the cortical hand territory in the primary somatosensory cortex (hereafter S1), suffers an extreme loss of sensory input in tandem with dramatic alterations of motor behaviour^1,2^. The functional and perceptual correlates of amputation-related plasticity are currently debated^3,4^. In particular, it is not clear whether functional cortical reorganisation is restricted to early life development or can also occur in adults.

Traditionally, research assessing cortical plasticity after upper-limb loss has followed the tenet that neighbouring body parts of the missing hand, and lower face in particular, shift and encroach into the deprived hand area. This emphasis on the lip representation stems from early electrophysiological work in non-human primates, where numerous studies demonstrated an ‘upside-down’ facial somatotopy, with the lower face immediately neighbouring the hand^5–13^. Here, the lips and/or lower-chin inputs have been shown to remap into the deprived hand area after sensory loss^14,15^, leading to the well-accepted assumption that remapping is determined by cortical proximity^16,17^. Thereafter, human measurement of topographic shifts has tended to focus on that of the lips, where researchers have reported that shifted lip representation towards and into the deprived hand area is significantly associated with phantom limb pain (PLP) intensity^18–22^. PLP is a neuropathic pain syndrome experienced in the missing, amputated limb by the majority of amputees^23^. This condition is commonly thought to arise from maladaptive cortical plasticity in S1 (although see^24^), specifically from a signal mismatch between the missing hand representation and the remapped inputs of the lips in the deprived hand area^25^.

The research focus on lip cortical remapping in amputees is based on this assumption that the lips neighbour the hand representation. However, only a handful of neuroimaging studies in humans has supported the inverted (or ‘upside-down’) somatotopic organisation of the face, similar to that of non-human primates^26,27^. Contrasting work alternatively suggests an upright orientation of the face in S1^28–32^, with the upper-face (i.e., forehead) bordering the hand area. In line with this, recent work reported that the shift in the lip representation towards the missing hand in amputees was minimal^33,34^, and likely to reside within the face area itself.

Surprisingly, there is currently no research that considers the representation of other facial parts, including the upper face in particular (e.g., the forehead), in relation to plasticity or PLP. Detailed mapping of the upper and lower face is therefore needed to assess typical topography of facial sensorimotor organisation, as well as remapping after limb loss.

Remapping after upper-limb loss has also been documented in individuals born without a hand (hereafter one-handers), who do not experience PLP^35^. Here it has been shown that the representation of multiple body parts, including the residual arm, legs and mouth, remapped into the missing hand territory^36,37^. Importantly, cortical remapping in this group does not depend on cortical proximity of the body parts. With regards to the lips, a recent transcranial magnetic stimulation (TMS) study has reported functionally-relevant lip activity in the deprived hand area of one-handers^38^, showcasing that reported remapping may also be functional. It was proposed that the observed remapping of various body parts could have been shaped by compensatory behaviour^36^, as these body parts are all used by one-handers to compensate for their missing hand function, but this hypothesis awaits validation.

Here, we conducted a mapping of face cortical organisation to determine facial orientation (upright versus inverted) in the primary sensorimotor cortex in 22 two-handed controls, 15 amputees and 21 one-handers. We used surface-based comparisons, including cortical (geodesic) distances, to measure the extent of cortical remapping of the upper (forehead) and lower face (lips) in relation to the deprived (or non-dominant) hand area across all groups. We also explored the representation of the tongue, which has not been previously studied in the context of deprivation-triggered brain plasticity. Furthermore, we used multivariate representational similarity analysis (RSA) in order to characterise more subtle alterations in the relationship between facial activity patterns (forehead, nose, lips, tongue) in the deprived hand and face areas, independent of gross spatial somatotopic shifts.

We found that facial topography was arranged in an upright manner, with the forehead (i.e., upper face) bordering the hand area across all groups. Contrary to traditional theories^39^, we did not find evidence for facial remapping, including the lips, into the deprived hand area of amputees. We did, however, observe significant remapping of all face parts (upper and lower face) in the one-handers’ group, validating our methodology as suitable for identifying remapping effects. Interestingly, remapping of the cortical neighbour (upper face) within the one-hander group was away from the missing hand area, while the lips and tongue representations shifted towards the deprived cortex, hinting that the underlying mechanism of remapping is more complex than simple cortical proximity.

## Materials and methods

### Participants

Seventeen individuals with acquired unilateral upper-limb amputation (age; *M*=53.71, *SE*=2.69, women; *n*=4, missing right hand; *n*=9), twenty-one individuals with unilateral congenital transverse arrest (age; *M*=42.67, *SE*=3.04, women; *n*=13, missing right hand; *n*=8) and twenty-two two-handed controls (age; *M*=45.55, *SE*=2.02, women; *n*=10, left-handers; *n*=6) were recruited (see Table 1 for full details). Two additional amputees who were recruited for the study did not participate in the scanning session due to MRI safety concerns, and further recruitment was stalled due to Covid-19 restrictions. The proportion of participants with intact/dominant right hand, as well as gender, were matched across groups (*χ*^2^_(2)_=2.674, *p*= .263; *χ*^2^_(2)_ = 5.593, *p* =.061)). While significant differences between groups were observed for age (*H*_(2)_=7.689, *p*=.021), post-hoc comparisons confirmed non-significant differences between amputees and one-handers relative to controls. Age covariates were therefore only included in statistical analyses when direct comparisons between amputees and one-handers were carried out. Procedures were in accordance with NHS National Research Ethics Service approval (18/LO/0474), and written informed consent was obtained.

**Table 1.**
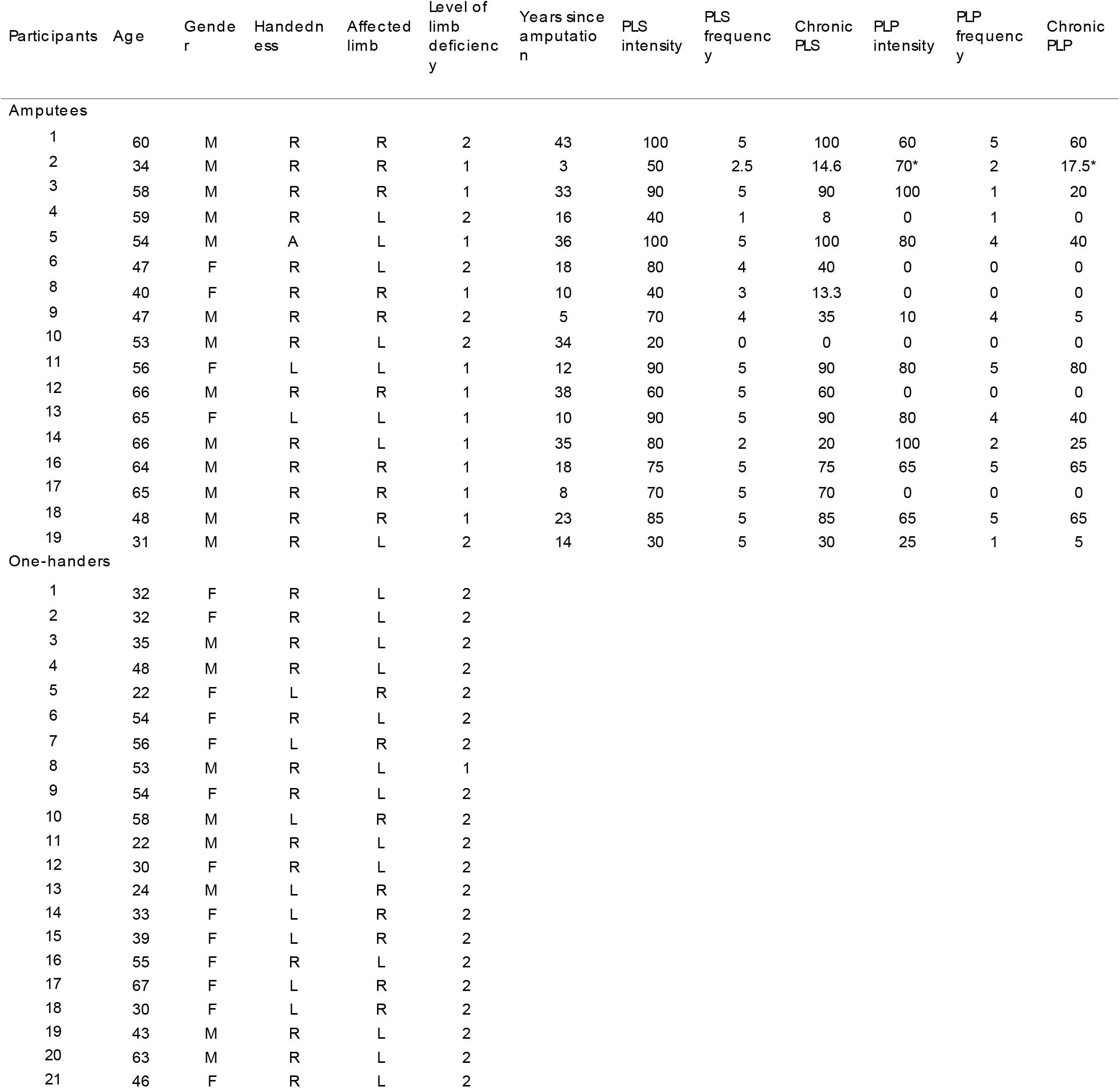
Demographic details for one-handed groups. Level of limb deficiency is as follows: 1 = limb loss above elbow (transhumeral), 2 = limb loss below elbow (transradial); L = left, R = right; PLS & PLP frequency: 0 = no sensation or pain, 1 = once or less per month, 2 = several times per month, 3 = once a week, 4 = daily, 5 = all the time. ^*^PLP intensity rating was on average. PLS = phantom limb sensations; PLP = phantom limb pain.

### Phantom sensations rating

Amputees were asked to rate the frequency of PLP experience within the last year. They also rated the intensity of their worst PLP experience during the last week (or in a typical week involving PLP; 0 = no pain, 100 = worst pain imaginable). A chronic measure of PLP was calculated by dividing the worst PLP intensity in the last week by PLP frequency (1 = all the time, 2 = daily, 3 = once a week, 4 = several times per month, 5 = once or less per month). This approach which takes into account the chronic aspect of PLP has been used successfully before ^33,35,40–44^, and has high inter-session reliability^42^. We also asked amputees about the vividness and frequency of non-painful phantom sensations (see Table 1).

### Functional MRI sensorimotor task

We used a facial active motor paradigm, where participants were visually instructed to move their forehead, nose, lips or tongue. This paradigm was chosen because it enabled bilateral activation of S1 simultaneously, allowing us to directly compare activity patterns between the two hemispheres (see Supplementary Figure 1 for validation of the active paradigm and Discussion for other considerations). Participants were also instructed to move their left and right thumb (amputees were asked to flex/extend their phantom thumb to the best of their ability; one-handers were asked to imagine such movement), resulting in 6 conditions. Baseline (i.e., rest) was included as a 7^th^ condition. Specific instructions involved: raising eyebrows (forehead), flaring nostrils (nose), puckering lips (lips), tapping tongue to the roof of the mouth (tongue), flexing and extending (thumb). The protocol comprised of 8 s blocks, with each condition repeated 4 times per run (5 times for baseline), over 3 functional runs. Before entering the scanner, participants practised each movement with the experimenter to ensure that the movement could be executed and to standardise each movement across participants (e.g., specificity and pace). Note that multiple participants reported during the experimenter briefing that they could not successfully flare their nostrils, and were therefore instructed to attempt moving their nose in the scanner.

### Functional MRI data acquisition and analysis

3T MRI data acquisition and pre-processing followed standard procedures, as detailed in the Supplementary Methods. Time-series statistical analysis was carried out using FMRIB’s Improved Linear Model (FILM). Task-based statistical parametric maps were computed by applying a voxel-based General Linear Model (GLM), as implemented in FEAT. The design was composed of 6 explanatory variables for each movement, convolved with a double-gamma hemodynamic response function^55^, and its temporal derivative. The six motion parameters were included as regressors of no interest. Motion outliers (> 0.9 mm) of large movements between volumes were included as additional regressors of no interest at the individual level (of total *n* volumes per group: amputees: 0.36%; controls: 0.36%; one-handers: 0.42%). For our main comparisons 6 contrasts were set up, corresponding to the facial movements’ (forehead, nose, lips, tongue and left/right thumb) relative to rest. Since the nose condition yielded weak activity relative to baseline, we excluded it from the univariate analysis (see Supplementary Figure 2).

The estimates from the three functional runs were then averaged voxel-wise using a fixed effects model in participants structural space, with a cluster forming z-threshold of 2.3 and family-wise error corrected cluster significance threshold of *p*< 0.05. Each estimates’ average was masked prior to cluster formation with a sensorimotor mask, defined as the precentral and postcentral gyrus from the Harvard Cortical Atlas. The sensorimotor mask was registered to the individuals structural scan using an inversion of the nonlinear registration by FNIRT. All functional MRI analysis was carried in individual’s native anatomical space.

### Regions of Interest (ROI) definition

Facial topography and remapping were studied using anatomical ROIs for the hand and face areas in S1. Although the primary motor cortex (M1) and S1 are expected to activate during facial movement we primarily focused on S1 remapping due to the traditional focus in the maladaptive plasticity literature on S1 representational shifts^39^. Furthermore, M1 topography tends to be less well-defined^49,50^, and so characterisation of typical facial topography may be more apparent in S1. Nevertheless, we wish to note that due to the proximity of S1 to M1, it is possible that marginal contribution from M1 may have affected our S1 activity profiles.

Firstly, S1 was defined on the average surface using probabilistic cytoarchitectonic maps, by selecting nodes for Brodmann areas (BAs) 1, 2, 3a and 3b^51^. The S1 hand ROI (hereafter hand ROI) was defined by selecting the nodes approximately ~1 cm below and ~2.5 cm above the anatomical hand knob. In contrast to earlier work^52^, we defined a conservative lateral boundary of the hand ROI (~1cm below the hand knob) to ensure there was limited facial activity captured. From the remaining parts of S1, the medial region was discarded and the lateral region was selected as the preliminary approximation for the S1 face ROI (hereafter face ROI; Fig. 1A).

**Figure 1.**
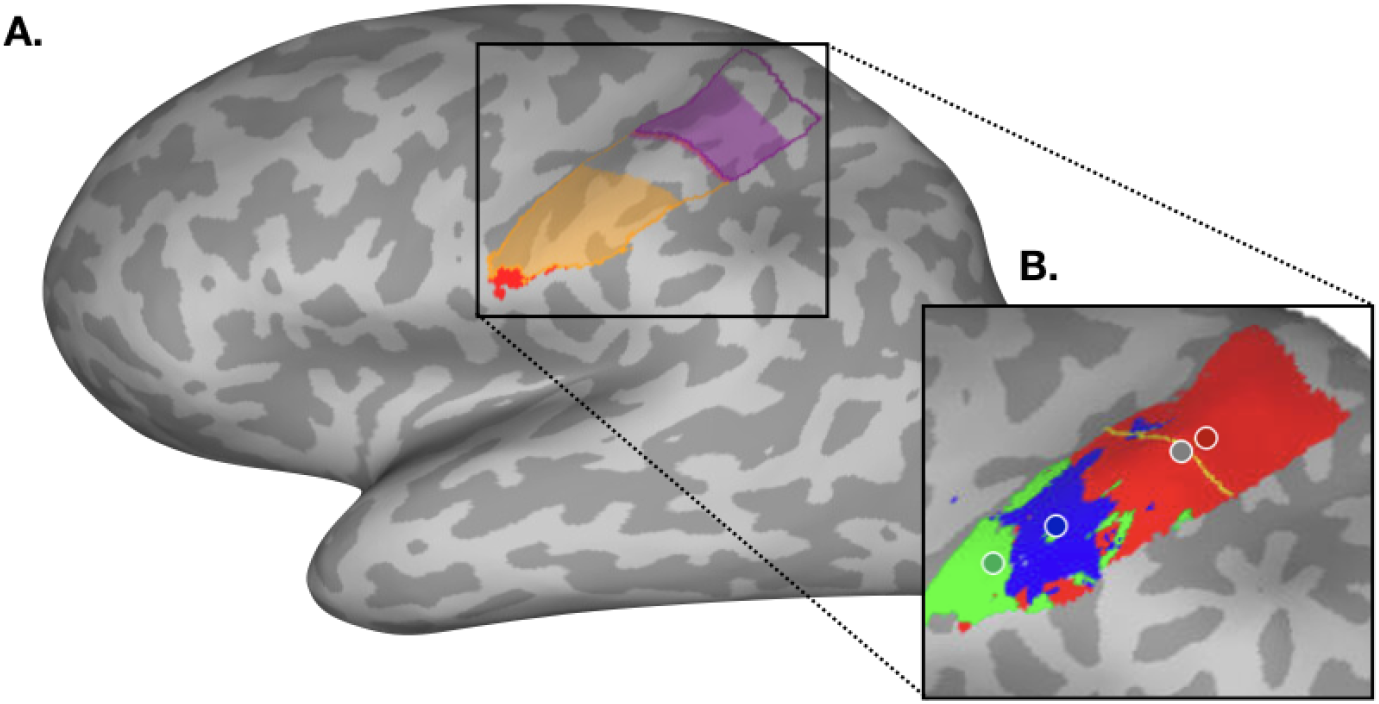
Regions of interest and winner-takes-all analysis in the primary somatosensory cortex for an example participant. (**A)** Regions of interest (ROI) used for univariate analyses are outlined in purple for the hand, and in orange for the face. Shaded areas of each region of interest denote the trimmed ROIs used for multivariate analyses. ROI overlap with the secondary somatosensory cortex (S2) is highlighted in red, and was removed from the face ROI in order to minimise somatotopic contribution from that region. (**B)** A typical winner-takes-all map from an example participant, with forehead activity in red, lip activity in blue and tongue activity in green. The centre-of-gravity for each movement is signified by a coloured dot outlined in white. The hand-face border is outlined in yellow, with the midpoint denoted by a grey dot. Cortical geodesic distances were measured from each facial parts CoG to the hand-face border midpoint.

Structural T1-weighted images were then used to reconstruct pial and white-grey matter surfaces using Freesurfer (version 7.1.1) at the individual level. The hand and preliminary face ROIs were then projected into individual brains via the reconstructed individual anatomical surfaces. As the secondary somatosensory cortex (S2) contains a crude somatotopy^53^, the preliminary face ROI was further trimmed in participant’s structural space by removing the overlap with S2. S2 was defined in MNI152 space using the Juelich Histological Atlas^54^. The S2 ROI was registered to participants’ structural space using an inversion of the nonlinear registration carried about by FNIRT. The remaining preliminary face ROI with the overlap from S2 removed was used as the face ROI for all univariate analyses. We note that due to the probabilistic nature of these masks, there could be some marginal contribution from S2 in our estimated face area.

### Winner-takes-all approach

To characterise S1 facial topography, the hand and face ROIs were combined to produce an overall S1 ROI (minus the medial region), and a winner-takes-all approach was used (Fig. 1B). For each participant, thresholded z-statistics averaged across the three functional runs were assigned to one of three face parts (forehead, lips, tongue), dependent on which facial movement relatively showed maximal activity within the S1 ROI. Face-winners (i.e., the output of the winner-takes-all) were then projected to the individual’s anatomical surface. Note that we excluded the thumb, which covered ~66% of the deprived hand ROI surface area in amputees and controls (see Supplementary Figure 3). This allowed us to align our analysis with previous research, and to draw comparisons of facial somatotopy across all groups (one-handers do not have a phantom limb, and therefore we cannot probe the ‘missing’ hand representation directly). All subsequent analyses at the individual’s anatomical surface level were computed using Connectome Workbench (v1.4.2).

### Cortical distance analysis

To assess possible shifts in facial representations towards the hand area, the centre-of-gravity (CoG, weighted by cluster size^33^) of each face-winner map was calculated in each hemisphere and the geodesic cortical distance between each movement’s CoG and a predefined cortical anchor computed. The cortical anchor was defined as the midpoint of the lateral border of the hand ROI (see Fig. 1B). The border was drawn manually and the midpoint calculated for each participant, and were visually confirmed by a second experimenter. The geodesic distance was assigned a negative value if the movement’s CoG was located below the hand border (i.e., laterally).

### Surface area calculation

To assess possible remapping into the hand area, a secondary winner-takes-all analysis was restricted to the hand ROI only. The surface area coverage (mm^2^) for each face-winner were computed on the individual anatomical inflated surface. We next calculated the proportion of the hand ROI occupied by each face part by dividing each face-winner’s surface area by the total hand ROI surface area for each individual. From the resulting percentages, we produced a laterality index for each movement with the following formula:

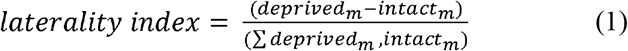

whereby *deprived*_*m*_ and *intact*_*m*_ represent the percentage of surface area coverage for the facial movement *m*, respectively in the deprived and intact hemisphere. A subsequent laterality index of +1 indicates surface area coverage of that movement solely within the deprived hemisphere (or the hemisphere contralateral to the non-dominant hand in controls), whereas a value of 0 represents an equal balance of surface area coverage across both hemispheres. Note that this approach characterises cortical remapping in relation to the intact hemisphere and has been used in numerous previous studies on amputees^18,19,21,22^. It assumes that the intact hemisphere reflects baseline (i.e., that it is truly ‘intact’), which may not be the case due to inter-hemisphere plasticity and/or homeostatic mechanisms^2,56,57^ and so also we compared our results to the control group.

### Multivariate representational analysis

Representational Similarity Analysis (RSA^62^) was used to assess the multivariate relationship between activity patterns generated by each face part. RSA was conducted in the hand and face ROIs to explore possible remapping across representational features between groups. To ensure the selectivity of the hand and face areas, the ROIs used for univariate analyses were each further trimmed medially by ~1cm, creating a 1cm gap between the hand and face ROIs. For each participant, parameter estimates for the four facial movements (forehead, nose, lips and tongue) and the contralateral thumb (for controls and amputees only) were extracted from all voxels within the chosen ROI, as well as residuals from each runs’ first-level analysis (three runs in total). Due to the increased sensitivity to subtle changes in activity patterns afforded by the multivariate approach, we included the nose movement. Multidimensional noise normalisation was used to increase reliability of distance estimates (noisier voxels are down-weighted), based on the voxel’s covariance matrix calculated from the GLM residuals. Dissimilarity between resulting facial activity patterns were then measured pairwise using cross-validated Mahalanobis distances^63^. Due to cross-validation, the expected value of the distance is zero if two patterns are not statistically different from each other. Distances significantly different from zero indicate the two representational patterns are different; negative distances indicate noise. Larger distances for movement pairs therefore suggest greater discriminative ability for the chosen ROI. The resulting six unique inter-facial representational distances (10 unique distances when including the thumb for controls and amputees only) were characterised in a representation dissimilarity matrix (RDM). Multidimensional scaling (MDS) was also used to project the higher-dimensional RDM into lower-dimensional space, whilst preserving inter-facial dissimilarity, for visualisation purposes only. Analysis was conducted on an adapted version of the RSA Toolbox in MATLAB^62^, customised for FSL^64^.

### Statistical analyses

All statistical analyses were carried out using JASP (Version 0.14). Outliers were classified as +/− 3 standard deviations to the mean. We chose to not remove outliers in our analyses and checked that the significance and direction of the results did not change if identified outliers were removed. When appropriate, univariate analyses was compared using parametric statistics. To assess normality for parametric tests, Shapiro-Wilk tests were run on residuals in combination with inspection of Q-Q plots and reporting of Levene’s Test for Equality of Variances. Where stated, non-parametric test statistics are reported where the assumption of normality has been violated. Analysis of Variance (ANOVA) was used to explore group differences to controls in cortical distances. Each mixed ANOVA had a between-subject factor of Group (Controls x Amputees; Controls x One-handers) and a repeated-measures factor of Hemisphere (Intact/Dominant x Deprived/Non-dominant), and was run separately for each facial movement (forehead, lips and tongue; see Supplementary Tables 1-2 for main effects). We controlled for brain size volume when comparing cortical distances between groups. Post-hoc comparisons were conducted with a Bonferroni correction for multiple comparisons (corrected alpha = .025; reported uncorrected *p*-values in text). If assumptions of normality were violated, the difference between cortical distances in the intact and deprived hemisphere were calculated, and the group difference between one-handed groups and controls was computed using a Mann-Whitney U test. Resulting statistics are reported alongside the mixed ANOVA output. Independent *t*-tests were used to calculate laterality indices group differences. We reported the corresponding Bayes Factor (*BF*_*10*_), defined as the relative support for the alternative hypothesis, for non-significant interactions and post-hoc comparisons. While it is generally agreed that it is difficult to establish a cut-off for what consists sufficient evidence, we used the threshold of *BF*<1/3 as positive evidence in support of the null, consistent with others in the field^65,66^ (though see ^67^). The Cauchy prior width was set at 0.707 (JASP’s default). To investigate whether remapping measures were related to PLP, we used a one-tailed Mann-Whitney U tests to compare laterality indices of amputees with (n=11) and without PLP (n=6) for relevant facial parts, under the hypothesis that PLP should result in greater remapping (see Supplementary Figure 4 for the analogous analysis for the geodesic distances).

For multivariate analyses, in order to quantify the dissimilarity, or ‘information’, each ROI holds for the face across groups, a linear mixed model (LMM) analysis was used, allowing us to consider the distinct contribution of face-thumb or face-face pairs. The face-face LMM contained fixed factors of group (controls, amputees and one-handers), hemisphere (intact, deprived) and facial pairs (6 unique representational distances). A random effect of participant, as well as covariates of age and gender, were also included in the model. An additional LMM was used to explore potential remapping in the hand ROI in relation to the non-dominant/phantom thumb in controls and amputees. Here the LMM contained the fixed factors of group (controls and amputees), hemisphere (intact, deprived) and face-to-thumb pairs (4 unique representational distances), as well as a random effect of participant (see Supplementary Tables 3-5 for main effects). All LMM’s were carried out in Jamovi (version 1.6.15) under restricted maximum likelihood (REML) conditions with Satterthwaite adjustment for the degrees of freedom.

### Data availability

Full data used for running the statistical analyses will be made available to the public upon publication via OSF. Data sharing will be provided prior to publication upon request.

## Results

### The cortical neighbour of the hand representation is the forehead

When looking at facial organisation at the group-level we found qualitatively similar activity maps across groups (see Fig. 2), highlighting a robust somatotopy of the face with preserved symmetry across the two hemispheres. These facial maps also indicate an upright orientation of the face in S1, with the forehead located closest to the hand area, followed by the lips, and the tongue located laterally, across all groups. The facial somatotopy presented here therefore suggests that the hand’s cortical neighbour is the forehead (or upper face), highlighting the need to reassess the often-cited, traditional lip-to-hand marker of cortical remapping in amputees and one-handers. However, conclusions based on group averages may be misleading as they ignore inter-individual differences.

**Figure 2.**
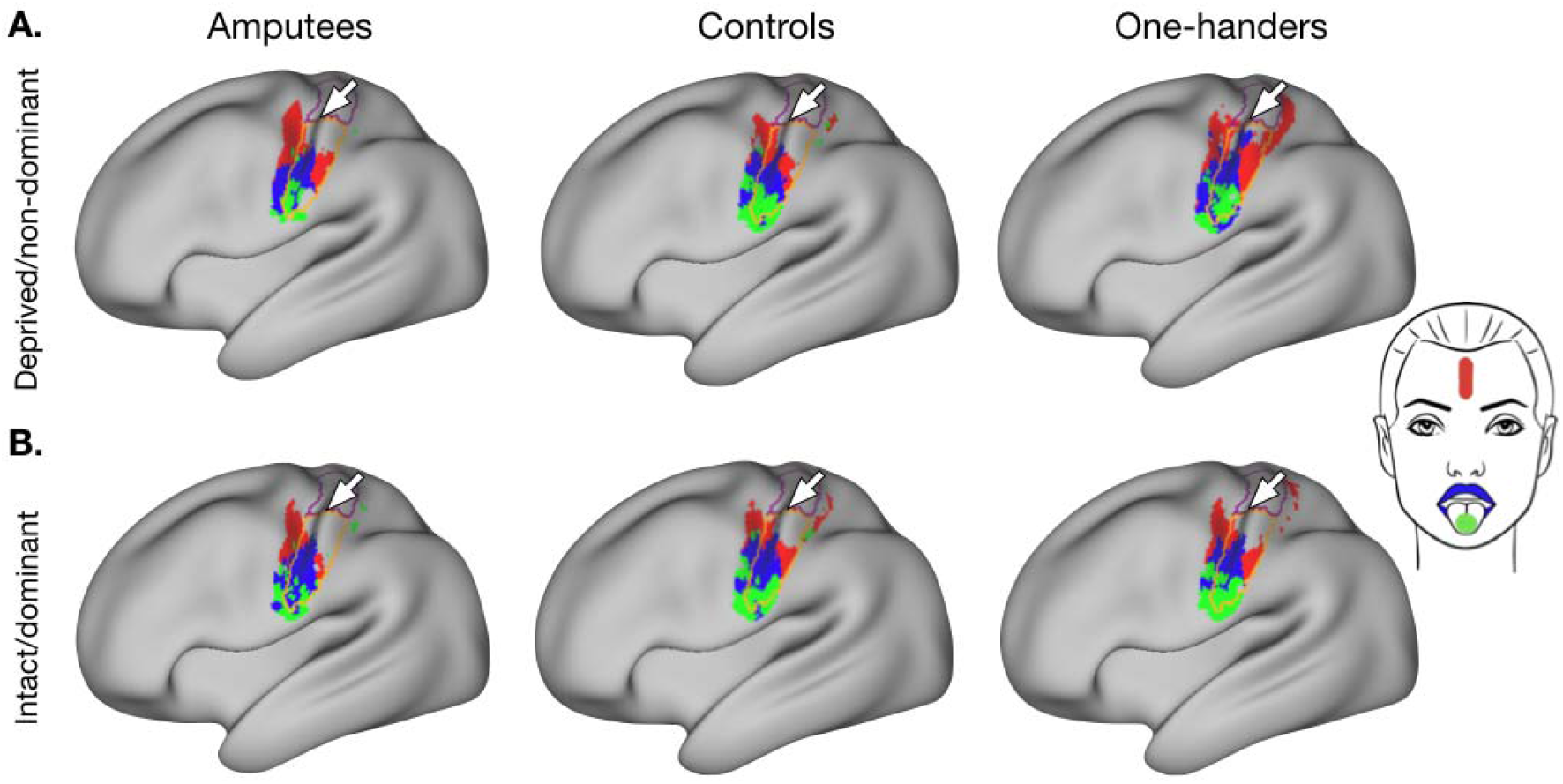
Group-level activity maps for each facial movement. Group average activity for the forehead (red), lips (blue) and tongue (green) movements, contrasted to rest, in the **(A)** deprived/non-dominant and **(B)** intact/dominant hemisphere for controls (n = 22), amputees (n = 17) and one-handers (n = 21). All clusters were created using a threshold-free cluster enhancement procedure with a sensorimotor pre-threshold mask (defined using the Harvard Cortical Atlas), and thresholded at p < .01. The hand and face ROIs are outlined in purple and orange respectively, and the central sulcus is denoted with a white arrow. Full methods for the construction of the group maps are available in the Supplementary Methods.

### One-handers, but not amputees, show lip remapping in the deprived cortex based on univariate topographic mapping

To account for inter-individual differences in functional topography and brain topology, we first explored changes in the cortical (geodesic) distance between the lips and the hand-face border of amputees and controls. Here we found no statistically significant main effects or group x hemisphere interaction (*F*_*(1,36)*_=0.003, *p*=0.954, *n*^*2*^=2.413_e-5_, *BF*_*10*_=0.449; controlled for brain size volume; Fig. 3B), indicating that the lip area in amputees is not located differently to that of controls. This is further supported by a lack of evidence for significant remapping of the lips (i.e., greater surface coverage) in the missing hand ROI for amputees when compared to controls (*U*=171.000, *p*=0.660, *d*=−0.086, *BF*_*10*_=0.355; Fig. 3C). These results suggest, contrary to popular theories on brain plasticity in amputees^39^, that the lips do not remap into the deprived hand area. However, the reported Bayes Factors indicated only anecdotal evidence for the null. We next compared the lips laterality index between those individuals who reported suffering from PLP (*n*=11) and those who no longer experienced chronic PLP (*n*=6) and found no significant differences (*U*=36.000, *p*=0.404, *d*=0.091, *BF*_*10*_=0.545).

When looking at lip plasticity within the one-handers group, however, we did note a slight qualitative shift in the location, and spread, of the lip activity within the deprived hemisphere (Fig. 2B). This is further supported by a visible shift of the one-handers lip group-level consistency map towards, and into, the deprived hand area (Fig. 3A). These changes in the lip representation were statistically significant, with a significant group x hemisphere interaction for the lips cortical distance to the hand-face border in one-handers and controls (*F*_*(1,40)*_=5.419, *p*=0.025, *n*^*2*^=0.032; controlling for brain size; Fig. 3B). Confirmatory comparisons indicated no statistically significant shifts of the lip CoG in the deprived hemisphere when compared to the controls non-dominant hemisphere (*t*_*(41)*_=−1.513, *p*=0.138, *d*=−0.462, *BF*_*10*_=0.745; Fig. 3B). However, shorter distances from the lips to the hand area were found in the deprived hemisphere of the one-handers when compared to their intact hemisphere (*t*_*(20)*_=−3.073, *p*=0.006, *d*=−0.671), indicating substantial evidence for lip remapping. These shifts in the deprived hemisphere were also reflected in significantly greater surface area coverage of the lips in the hand ROI when compared to controls (*U*=119.000, *p*=0.007, *d*=−0.485; Fig. 3C), which was significantly different from zero (*W*=197.000, *p*=0.003, *d*=0.706). This evidence of lip remapping is in line with previous work in one-handers^36,38^.

**Figure 3.**
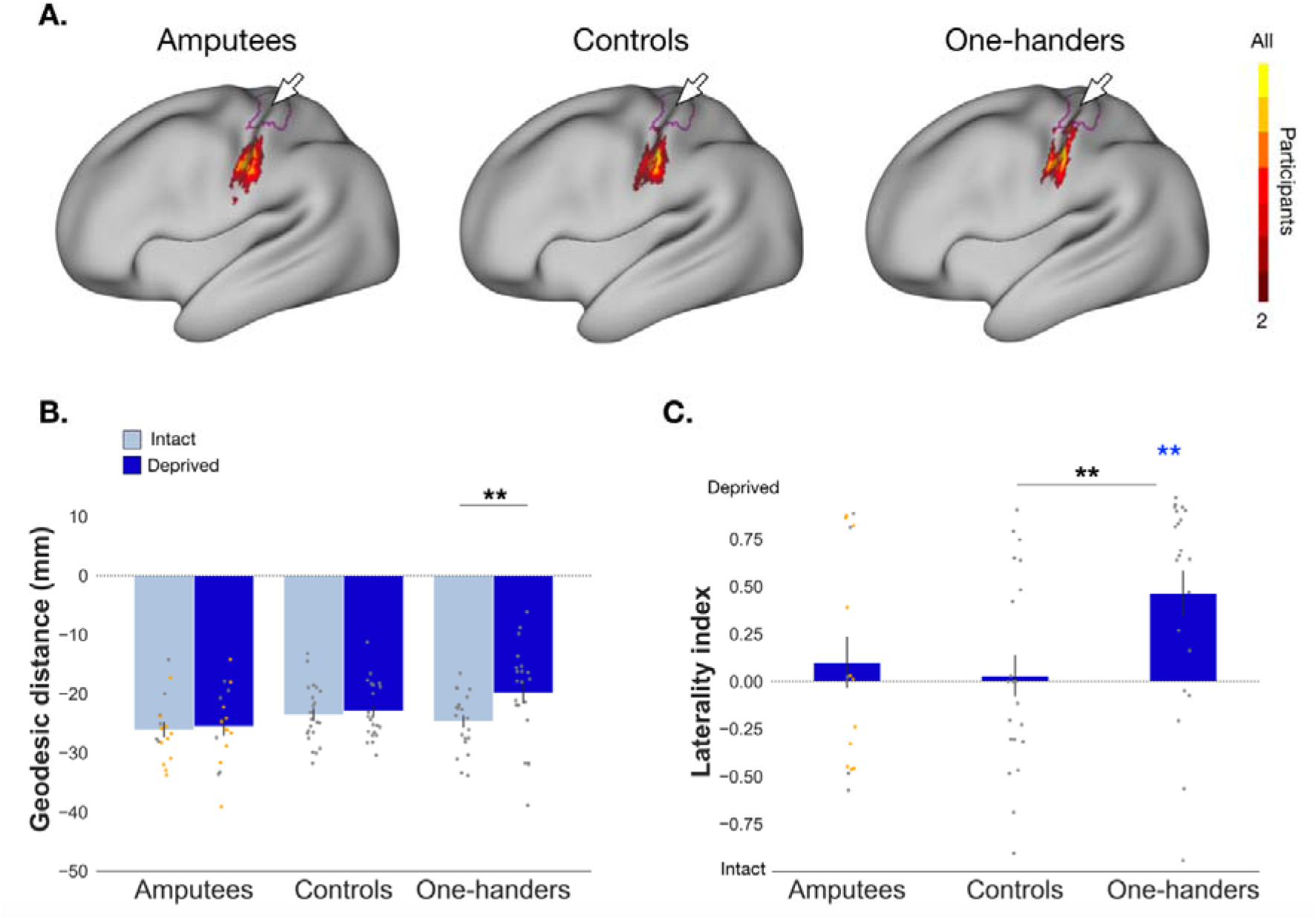
Characterisation of lip (re)mapping in the primary somatosensory cortex. **(A)** Group-level consistency map for the lips in the S1 ROI (hand and face combined) across groups. The colour gradient represents participant agreement for maximally activating that particular voxel, relative to other face movements (winner-takes-all approach; See the Supplementary Methods for full details on the construction of these maps). The hand ROI is outlined in purple and central sulcus denoted by the white arrow. **(B)** Cortical geodesic distances from the lip CoG to the hand-face border are plotted for amputees (n = 17), controls (n = 22) and one-handers (n = 21). Distances in the intact/dominant hemisphere are plotted in light blue, and distances in the deprived/non-dominant hemisphere are plotted in darker blue. Positive distances indicate the lips CoG is located medial to the hand-face border in the hand ROI, and negative distances indicate the lips CoG is located lateral the hand-face border in the face ROI. The hand-face border itself equates to a geodesic distance of zero. **(C)** Laterality indices for the proportion of surface area coverage of the lips in the hand ROI for all groups (amputees, controls and one-handers). Positive values indicate greater surface area coverage in the deprived/non-dominant hemisphere, and negative values reflect greater surface area coverage in the intact/dominant hemisphere. Standard error bars and all individual data-points are plotted in grey and uncorrected for brain size. Amputees with PLP (yes/no) are plotted in orange. * p < .05; ** p < .01 (please note p-values are uncorrected); coloured asterisk’s indicate values are significantly different from zero.

### One-handers, but not amputees, show forehead remapping in the deprived cortex, based on univariate topographic mapping

As we note a qualitative upright orientation of the face (see Fig. 2), the question remains as to whether the neighbour to the hand – the forehead – would reorganise after limb loss in amputees, as hypothesised by traditional theories^39^. Again, we found no significant evidence for cortical remapping of the neighbouring forehead in amputees when assessing changes in cortical distances (group x hemisphere: *F*_*(1,36)*_=0.935, *p*=0.340, *n*^*2*^=0.011, *BF*_*10*_=0.138; controlled for brain size volume; non-parametric equivalent: *U*=231.000, *p*=0.221, *d*=0.235, *BF*_*10*_=0.498; Fig. 4B). A significant difference was found, however, of reduced forehead surface area coverage in the deprived hand ROI when compared to controls (*t*_*(37)*_=2.048, *p*=0.048, *d*=0.661, *BF*_*10*_=1.572; Fig. 4C). Interestingly, the direction of results indicates less, not more, remapping of the forehead in the deprived hand ROI of amputees. Despite this, we found non-significant differences when comparing the forehead laterality index for amputees with and without PLP (*U*=26.000, *p*=0.769, *d*=−0.212, *BF*_*10*_=0.322). Taken together, these results suggest that if remapping of the cortical neighbour – the forehead – does occur, this effect is small in comparison to controls, away from the hand area, and is not related to PLP.

**Figure 4.**
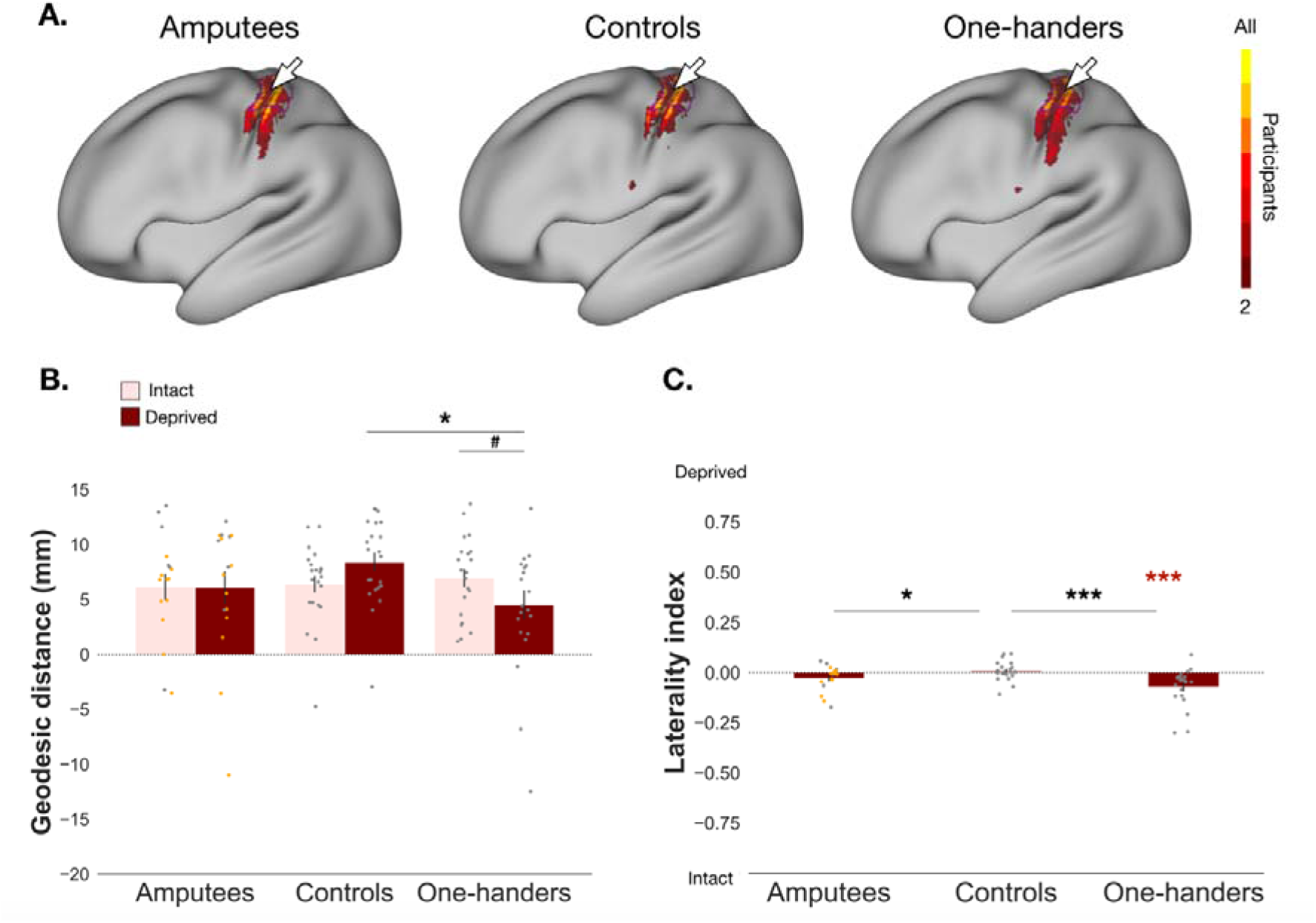
Forehead topography in the primary somatosensory cortex. All annotations are as in Figure 2. Distances in the intact hemisphere are plotted in pink and deprived hemisphere in red. **^#^** strong positive trend; * p < .05; *** p < .001; coloured asterisk’s indicate values are significantly different from zero.

When looking at the one-handers group we did find significant evidence of forehead remapping with a group x hemisphere interaction (*F*_*(1,40)*_=8.287, *p*=0.006, *n*^*2*^=0.059; controlled for brain size volume; non-parametric equivalent: *U*=343.000, *p*=0.006, *d*=0.485; Fig. 4B). Confirmatory comparisons indicated a positive trend for shorter distances of the foreheads’ CoG to the hand-face border in the deprived hemisphere when compared to their intact hemisphere (*t*_*(20)*_=2.349, *p*=0.029, *d*=0.513, *BF*_*10*_=2.094) and significantly shorter distances when compared to the controls non-dominant hemisphere (*U*=332.000, *p*=0.014, *d*=0.437). As the forehead’s CoG tended to be located above the hand-face border (see Fig. 4A), these results indicate a significant shift of forehead activity away from the deprived hand ROI. This is further supported by a significant decrease of surface area coverage for the forehead in the deprived hand ROI when compared to controls (*U*=381.000, *p*< .001, *d*=1.069), which was significantly different from zero (*W*=19.000, *p*< .001, *d*=−0.835; Fig. 4C). Remapping of the cortical neighbour in one-handers, therefore, manifests in a shifting away of the upper face from the deprived hand area, possibly due to increases in activity of other facial movements, e.g., lips.

### Tongue movements produce different topographic maps across groups

We also assessed changes in the tongue representation, which is not an immediate neighbour to the hand in S1(Fig. 5A). We did not find significant evidence for shifts in the tongue’s CoG towards the hand-face border in amputees when compared to controls (group x hemisphere: *F*_*(1,36)*_=2.546, *p*=0.119, *n*^*2*^=0.007, *BF*_*10*_=0.182; controlled for brain size volume; non-parametric alternative: *U*=126.000, *p*=0.087, *d*=−0.326, *BF*_*10*_=0.882; Fig. 5B). Nevertheless, the tongue did show significantly greater surface area coverage in the deprived hand ROI of amputees when compared to controls (*t*_*(37)*_=−2.759, *p*=0.009, *d*=−0.891; Fig. 5C), which was also significantly different to zero (*t*_*(16)*_=2.302, *p*=0.035, *d*=0.558). As tongue remapping is not reflected consistently across analyses, and due to the lack of pre-existing hypotheses, this preliminary result should be interpreted with caution. However, it does indicate that some level of cortical remapping may occur in amputees after limb loss.

**Figure 5.**
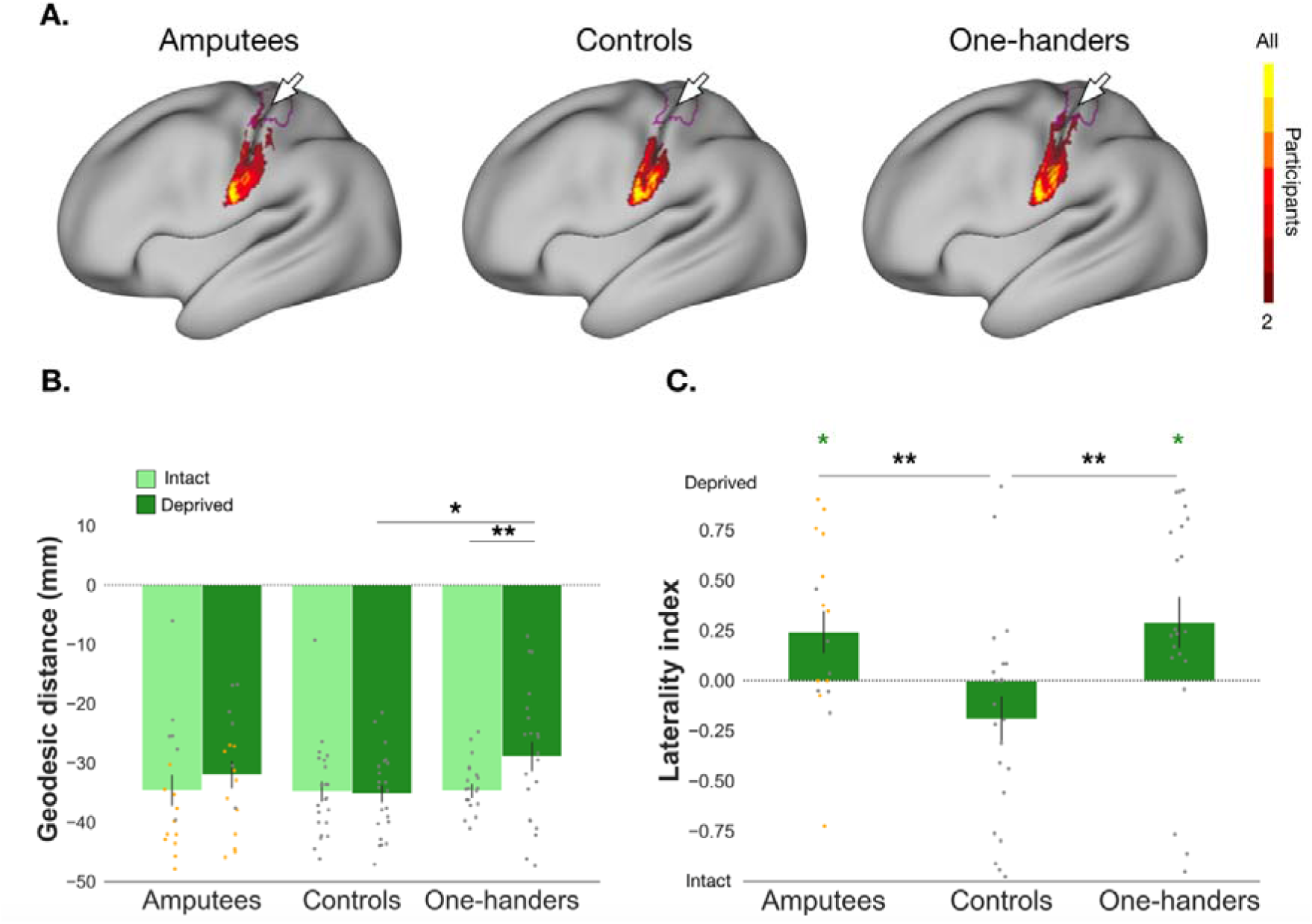
Tongue remapping in amputees and one-handers in the primary somatosensory cortex. Distances in the intact hemisphere are plotted in light green and distances in the deprived hemisphere in dark green. All other annotations are as in Figure 2.

We next explored whether this increase in tongue activity within the deprived hand ROI, as captured by the laterality index of amputees, was related to PLP (Fig. 5C), and found a non-significant difference (*t*_*(15)*_=1.223, *p*=0.120, *d*=0.620, *BF*_*10*_=1.169). These results suggest, along with an inconclusive Bayes Factor, that amputees with PLP do not report greater instances of tongue remapping, when compared to amputees without PLP.

We also found greater surface area coverage of the tongue in the deprived hand ROI of one-handers (see Fig. 5A) that was significantly different from zero (*W*=180.000, *p*=0.024, *d*=0.558) and from controls (*U*=109.000, p=0.003, *d*=−0.528; Fig. 5C). This increased tongue activity in the one-handers deprived hand area was also reflected in a significant group x hemisphere interaction (*F*_*(1,40)*_=7.149, *p*=0.011, *n*^*2*^=0.032; controlled for brain size volume; non-parametric equivalent: *U*=140.000, *p*=0.027, *d*=−0.394 Fig. 5B) of the cortical distance between the tongues’ CoG and the hand-face border. Confirmatory comparisons indicated significantly shorter distances to the hand-face border for the tongue in the deprived hemisphere compared to intact (*t*_*(20)*_=−2.902, *p*=0.009, *d*=−0.633), and when compared to the controls’ non-dominant hemisphere (*U*=150.00, *p*=0.050, *d*=−0.351). These results tentatively suggest that cortical remapping in one-handers may extend further to include tongue facial movements.

### Brain decoding in the deprived hand area reveals stable facial representational pattern for amputees, and increased facial information in one-handers

The analyses described above focused on the topographic relationship of the three facial parts, but cortical remapping could potentially manifest subtly, without disrupting the spatial distribution of the face representation. RSA identifies statistical (dis)similarities across activity patterns, providing a more sensitive measure of representational changes^68^. To further explore any possible remapping in the deprived hand area of the amputees, we first looked at the pattern of dissimilarity for each face part relative to non-dominant/phantom thumb in the hand ROI for controls and amputees. Here we found a non-significant group x hemisphere x face-thumb interaction (*F*_*(3,259.0)*_=0.569, *p*=0.636), as well as a non-significant group x hemisphere interaction (*F*_*(1,259.0)*_=0.057, *p*=0.811; Fig. 6A). The pattern of facial activity in relation to the thumb is therefore statistically comparable between amputees and controls, indicating similar representational structure of the face, relative to the hand, between the two hemispheres and groups.

**Figure 6.**
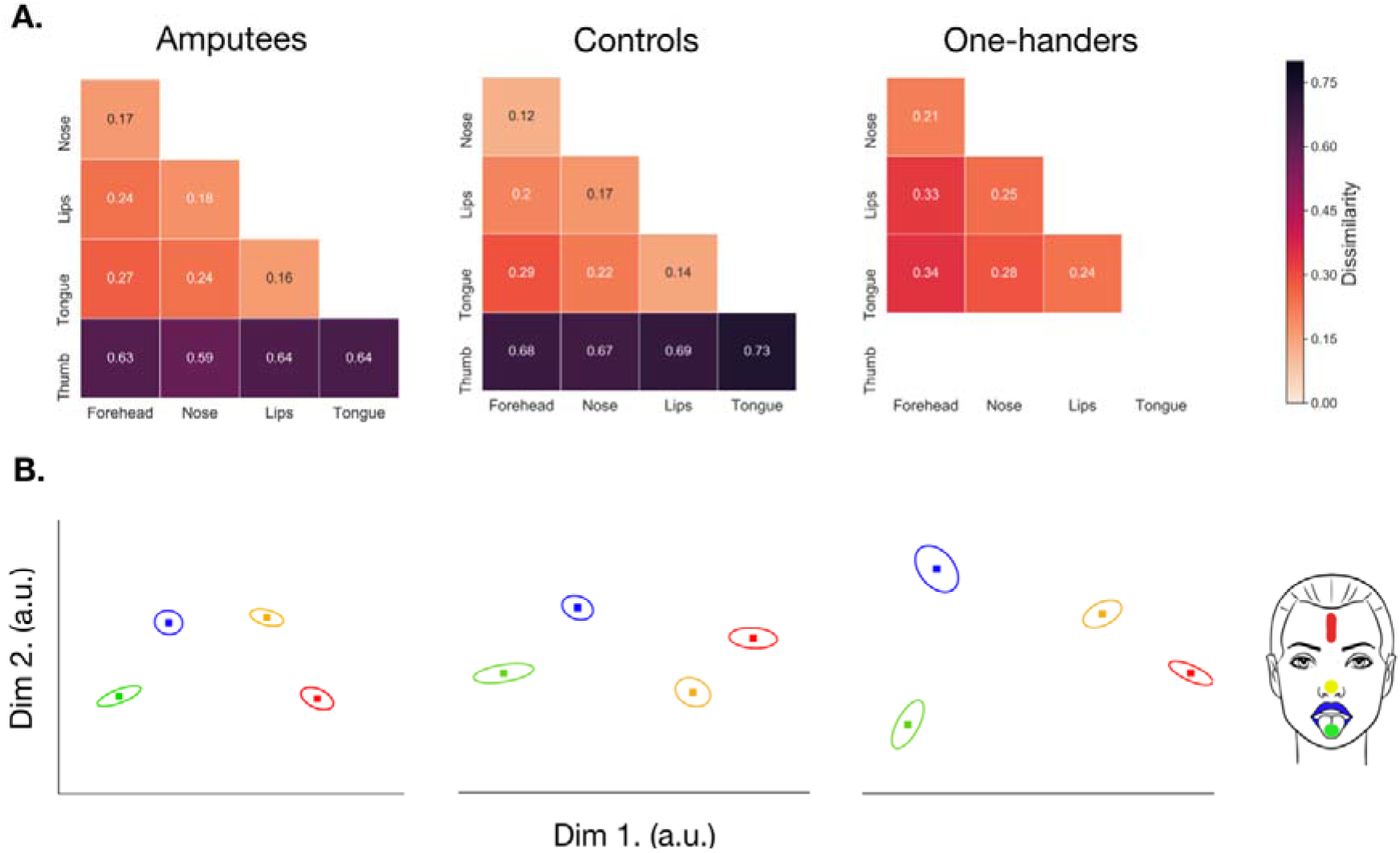
Representational Similarity Analysis (RSA) in the deprived/non-dominant hand area across all groups. **(A)** Representational Dissimilarity Matrices (RDMs) for amputees (n = 17), controls (n = 22) and one-handers (n =22). Greater dissimilarity between activity patterns for the chosen pairwise comparison indicates more information for that facial part within the hand area. Smaller dissimilarity values of facial (and thumb) activity patterns indicates a reduced ability to discriminate between the chosen movements in the hand area. Face-thumb distance values are only shown for the controls and amputees. (**B)** Multi-dimensional scaling plots for each group, which projects the RDM distances into a lower-dimensional space. Here the distances between each marker reflects the dissimilarity, with more similar activity patterns represented closer together, and more distinct activity patterns positioned further away. Forehead movements are plotted in red, with the nose in yellow, lips blue and tongue green, and the standard error is plotted around each data point. Please note, a different scale was used compared to the face ROI (Figure 7).

**Figure 7.**
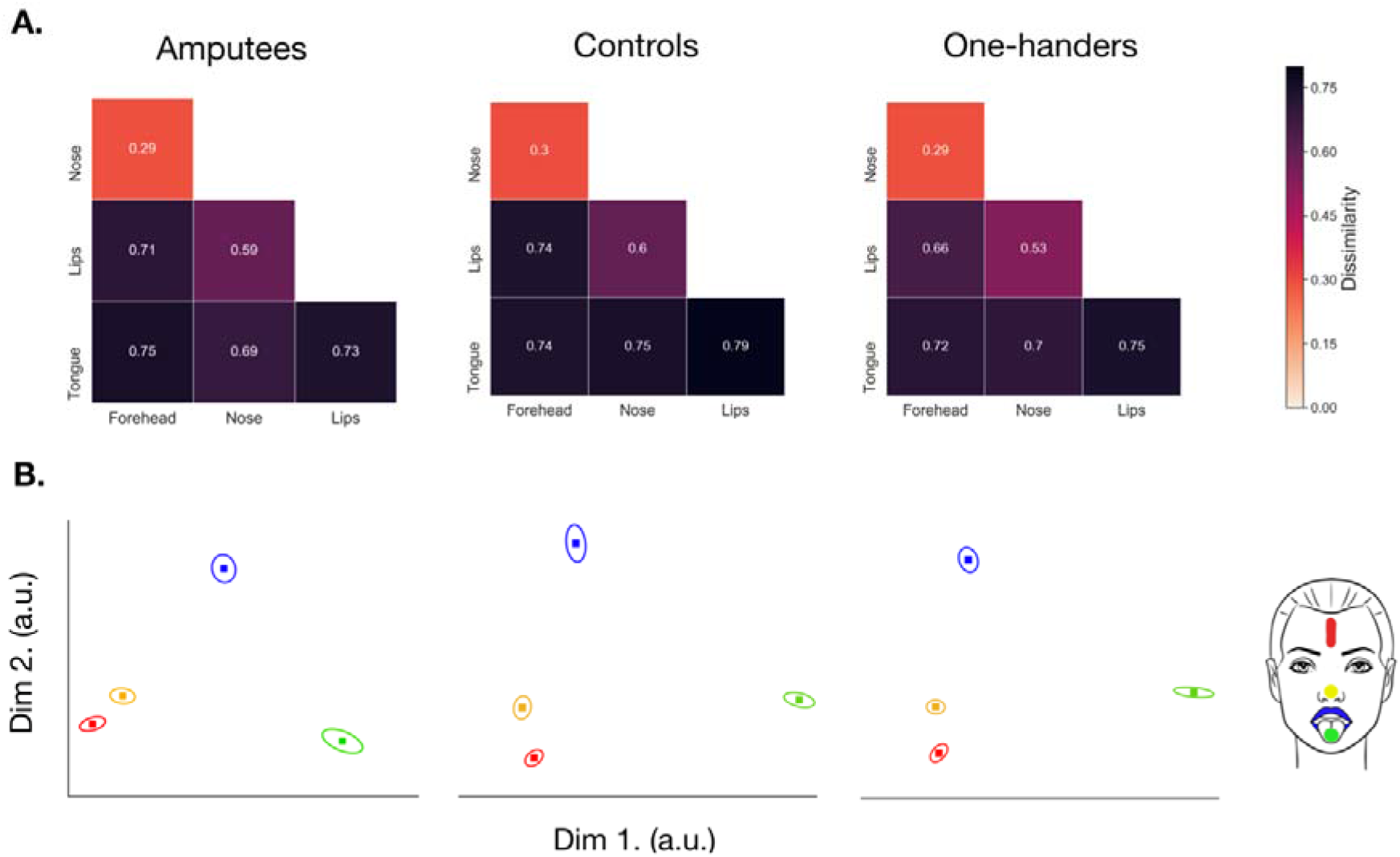
Representational Similarity Analysis (RSA) in the deprived/non-dominant face area across all groups. All annotations are as in Figure 6.

We also found a non-significant group x hemisphere x face-face interaction when looking at face-face pairwise dissimilarity in the hand ROI across all three groups (*F*_*(10,627.0)*_=0.572, *p*=0.837; controlled for age; Fig. 6B), suggesting a similar representational structure of the face across hemispheres and groups. However, when we looked at the average amount of facial information within the hand ROI, we did find a significant group x hemisphere interaction (*F*_*(2,627.0)*_=14.544, *p*< .001), indicating potential differences in facial information content in the hand area. Post-hoc comparisons (corrected alpha=0.0125; uncorrected *p*-values reported) exploring this effect reported significantly greater dissimilarity between facial-part representations in the deprived hemisphere of the amputees (*M*=0.214; *SE*=0.021; t_(627.0)_=−4.401, *p*< .001) and one-handers (*M*=0.273; *SE*=0.018; t_(627.0)_=−5.668, *p*< .001), when compared to their respective intact hemisphere (amputees: *M*=0.152; *SE*= 0.0205; one-handers: *M*=0.202; *SE*=0.018). When comparing to the controls non-dominant hemisphere, we only found significantly greater facial information in the one-hander’s deprived hand area (t_(72.8)_=−3.297, *p*=0.002) and a non-significant effect for amputees (t_(71.7)_=−0.828, *p*=0.411). These results are in line with our univariate analyses, which demonstrate significant cortical remapping of facial parts in the one-handers group. It also suggests that there may be inter-hemispheric changes in facial information in the intact hand ROI of amputees (*M*=0.152, *SE*=0.021; controls: *M*=0.207, *SE*=0.017), though this latter result awaits further confirmation.

For completion, we also looked at facial activity patterns (i.e., face-face pairwise dissimilarities) within the face ROI across all three groups. Here we found non-significant differences for a group x hemisphere x face-face interaction (*F*_*(10,627.0)*_=0.136, *p*=0.999) and group x hemisphere (*F*_*(2,627.0)*_=0.626, *p*=0.535), suggesting a similar representational pattern of facial activity, i.e., facial information content, across hemispheres and groups (Fig. 7).

## Discussion

It is a well-accepted notion, rooted in non-human primate electrophysiological data, that upper-limb amputation triggers cortical remapping of the assumed neighbour – the lower face – into the missing hand area. This previous work predominantly characterised remapping by investigating shifts of the lip representation^18–22,69,70^. However, by focusing on only one face part, activity elicited by other facial parts (such as the forehead) in areas of interest in S1 (i.e., the hand-face border) are not taken into account (see^2^). Here we explored the relationship of face-to-hand remapping in controls and one-handed groups, and used both univariate (topographic) and multivariate (representational structure) methods to investigate in detail the information content of the face in both the deprived and intact hand and face areas. We found evidence for an upright somatotopy of the face across all groups, suggesting that the cortical neighbour to the hand in humans is the upper, not lower, face. We further found little evidence for remapping of all tested facial parts in amputees, with no significant relationship to the presence of PLP. As a positive control, we also recruited individuals that were born without a hand (one-handers), who have previously shown cortical remapping across multiple body parts^36–38^. Across all facial parts, one-handers showed evidence for a complex pattern of face remapping in the deprived hand area, with consistent and converging evidence across analysis approaches. Together, our findings demonstrate that the face representation in humans is highly plastic, but that this plasticity is restricted by the developmental stage of input deprivation, rather than cortical proximity.

Firstly, our univariate analyses at both group and individual level confirmed an upright orientation of the face in controls, amputees and one-handers. Here we found that the forehead representation borders the hand area, followed by the lips, and tongue located more laterally. These results provide the first neuroimaging evidence of an upright representation of the face in humans. Indeed, while early and recent intracortical recordings and stimulation studies^28,29,31,71^ emphasised an upright organisation, in agreement with our results, neuroimaging studies in humans provided contradictory evidence for the past 30 years. For instance, early studies reported an inverted facial somatotopy^27,72^, similar to the topography reported in primates^11,12^. Many subsequent studies then found no clear somatotopic organisation^73–75^ or suggested an onion-like segmental topography^76^. These discrepancies and inconsistent neuroimaging findings may arise from the low-resolution of the techniques, small sample sizes and the challenge to find a robust and reliable method to stimulate face parts (and thus elicit detectable cortical activation). Our results provide converging evidence validating the upright orientation of the face, and indicate that the cortical neighbour to the hand is likely to be the upper face, which has important implications for cortical remapping theories based on cortical proximity^39^. If cortical remapping of neighbours exists, we would therefore expect to see the forehead shifting towards and into the hand area – not the lips.

It may be argued that it is difficult to achieve isolated execution of specific facial muscles when performing gross movements without impacting sensory processing of neighbouring facial parts. For instance, tongue movements in our paradigm (e.g., touching the roof of the mouth with the tongue), may be best considered as a holistic inner mouth movement, and forehead movements may be best considered as engaging the upper face. While this limitation is valid, it may also be relevant (though to a smaller degree) for passive paradigms, as stimulation can induce waves that propagate through the skin^77–79^ and Pacinian receptors were found to activate during stimulation of remote sites^80,81^. Despite this caveat, both our univariate and multivariate analyses showed that we were successful in isolating sensorimotor representations of the various movements (forehead, nose, lips and tongue) within our regions of interest (see Supplementary Figure 1 for validation). In other words, even if somatosensory information is overlapping across movements, there is still enough distinct information to separate representational patterns. This finding indicates the suitability of our motor paradigm for teasing apart facial somatotopy, allowing us to characterise the face in greater detail than previously attempted.

When looking at face-to-hand remapping in amputees, where the remapping of cortical neighbours has been the prevalent explanation for PLP, we find little evidence of shifts of locality and remapping in the deprived hand area for facial parts, including the neighbour (forehead) and hypothesised neighbour (lips). Our univariate results are further supported by our multivariate analysis, where we find no significant changes in the relationship between face-to-thumb activity in the deprived hand area. These results support previous work reporting a lack of cortical remapping after amputation^35,42,82^, suggesting that in amputees this area might be functionally unipotent – pertaining to hand-related activity alone and lacking the ability to rescope after hand loss. However, due to the inconclusive Bayes Factor in our key analyses, we cannot strongly conclude that remapping does not occur in this group. This could be attributed to our relatively small sample (further recruitment was prevented due to Covid-19 restrictions), and in particular, the small proportion of amputees experiencing PLP (11 out of 17). However, pain is not a necessary condition for deprivation-triggered remapping^83–85^ and vice versa, PLP can be experienced in absence of remapping^86^. Moreover, previous studies reporting significant difference between amputees who experienced PLP and those who do not, often employed a similar sample size^20^, indicating that the expected effect of remapping should be substantial.

We did find anecdotal evidence for remapping for the tongue within the deprived hand area in amputees. This was a surprising result, as the tongue is not a cortical neighbour to the hand, and was not specifically hypothesised to remap in amputees. We also found that amputees demonstrated a different amount of facial information across the two hand areas. Although this multivariate result was not significantly different to that of controls, it demonstrates the plausibility that cortical remapping in amputees may exist to a certain degree (e.g., an inter-hemisphere imbalance), albeit any relationship to PLP is tenuous. While these latter results require further validation, they support our premise that cortical proximity of representations may not be a necessity for remapping to occur. In this context, as our tongue condition could also be classed as an ‘inner mouth’ movement, it is important to note that previous work addressing sensorimotor representations of the mouth and the larynx have demonstrated both lateral and more medial ‘hotspots’ (i.e., a ‘double’ representation^87^). The potential tongue remapping in amputees, therefore, may reflect changes in the medial mouth representation, but this would need to be investigated further by separating the relative contributions of the tongue and larynx.

We did find converging and conclusive evidence for cortical remapping of all facial parts (neighbours and non-neighbours) in our (congenital) one-hander group. Here the pattern of remapping is strikingly different to that of cortical neighbourhood theories. Specifically, the location of the cortical neighbour – the forehead – is shown to shift away from the deprived hand area, which is subsequently ‘taken over’ by the lips and the tongue. The increase of facial activity in the deprived hand area is in turn supported by our multivariate results, whereby significantly greater information content for the face was found in the deprived hand area for one-handers when compared to controls. One-handers’ deprived hand area, therefore, seems to have increased discriminability between different facial movements. It is difficult to ascertain from our study the drivers of this remapping. It has been suggested previously that remapping within this group may be driven by functionally-relevant behaviour substituting the loss of the limb^36,38^. Alternative explanations relate to an overall and unspecific release of inhibition (i.e., decreased GABA) in the missing hand area, allowing for latent activity of other body parts to be detected^36^. While speculative, our results tend to support the former, as we report remapping for facial parts which have the ability to compensate for hand function, e.g., using the lips and/or mouth to manipulate an object, and a lack of remapping for those that cannot (the forehead). This increased activity from body parts compensating for hand function may represent a stabilising mechanism, aimed at preserving the integrity of the sensorimotor network and its function^2^. The deprived hand area in one-handers, therefore, may be deemed pluripotent – suitable for adapting to multiple body parts^38^, which may preserve the role of the hand area by sustaining its hand-function related information content.

A limitation that should be acknowledged arises from the potential contribution to S1 from M1 activity. Since these cortical areas are neighbours, it is difficult to separate them with certainty. We minimised the contribution of M1 by taking multiple acquisition and pre-processing steps, including the use of anatomical delineation at the individual level, as well as a comprehensive analytical approach (e.g., both univariate and multivariate techniques). Across the board we find robust evidence for remapping in congenital one-handers and no reliable evidence for similar remapping in amputees. Furthermore, it has been claimed that active movements may produce different cortical maps to those with passive stimulation^88,89^, and previous work demonstrating a relationship between cortical remapping and PLP tended to use passive stimulation^18,19,22,69^. However, we do not think this methodological difference underlies our contrasting results as movement-induced lip activity has been shown to demonstrate lip remapping before^20,70,90^, indicating that an active paradigm is suitable for demonstrating cortical remapping (if it exists). Conversely, a recent study using passive lip stimulation in amputees did not find any evidence for remapping^91^. Moreover, we recently ran a study which found that S1 topography and multivariate representational structure are similar across active and passive paradigms^92^. Moreover, the choice of an active paradigm is the most reflective of naturalistic tactile inputs in everyday life. Together with robust evidence for remapping in one-handers using all the methods tested here, our choice of active paradigm is clearly suitable to identify topographic organisation and remapping, and is practically accessible and translatable to fMRI designs.

Our findings seem contradictory to the many previous studies reporting lip remapping in amputees^18–22,69,70^. A major difference with regards to these previous studies, which predominantly focused on a single part of the face, lies in the fact that our study was the first to assess the mapping and potential remapping of multiple facial parts at once. By focusing on the lips only, previous designs excluded other facial parts which may have elicited greater activity in certain areas of S1, resulting in a less accurate delineation of the lip-selective representation. Such down-sampling of body maps, therefore, can lead to biased results and interpretation^2^. While our design is not exempt from this limitation, the fact that we assessed other parts of the face may explain why our results diverged from previous findings.

To conclude, our use of both univariate and multivariate analyses found consistent evidence for a complex pattern of face remapping in congenital one-handers, in line with the theory suggesting remapping in this group reflects compensatory behaviour^2^. This is in contrast to amputees, where we find little evidence for cortical remapping, indicating a stability of both the hand and face representation after limb loss. By and large, remapping measures were not linked to PLP. Our results call for a reassessment of traditional remapping theories based on cortical proximity, and future research into potential remapping of the inner mouth representation after limb loss.

## Abbreviations

Analysis of Variance: ANOVA
Bayes Factor: BF_10_
Brodmann areas: BAs
centre-of-gravity: CoG
fMRI: functional MRI
general linear model: GLM
linear mixed model: LMM
multidimensional scaling: MDS
PLP: phantom limb pain
PLS: phantom limb sensations
primary motor cortex: M1
primary somatosensory cortex: S1
representation dissimilarity matrix: RDM
representational similarity analysis: RSA
regions of interest: ROI
transcranial magnetic stimulation: TMS

## Acknowledgements

We thank Arabella Bouzigues and Maria Kromm for their substantial help in terms of recruitment and data collection, we also thank Adriana Zainurin, Esther Teo, Christine Tan, Raffaele Tucciarelli and Mathew Kollamkulam for help with data collection. We thank Opcare for their help with participants recruitment, and our participants and their families for their ongoing support to our research.

