## Supplementary Figure 1 for "Reassessing face topography in primary somatosensory cortex and remapping following hand loss"

**Supplementary Methods**

**MRI data acquisition**

Functional and anatomical MRI data were obtained using a 3 Tesla Prisma MRI scanner (Siemens, Erlangen, Germany) with a 32-channel head coil. Anatomical data was acquired using a T1-weighted sequence (MPRAGE), with the following parameters: TR = 2530 ms; TE = 3.34 ms; flip angle = 7°; voxel size = 1 mm isotropic resolution. Functional data based on the blood oxygenation level dependant (BOLD) signal were acquired using a multiband T2*-weighted pulse sequence, with a between-slice acceleration factor of 4 and no in-slice acceleration (TR = 1450 ms; TE = 35 ms; flip angle = 70°; voxel size = 2 mm isotropic resolution; imaging matrix = 106 x 106; FOV = 212 mm). 72 slices were oriented in the transversal plane. A total of 172 whole-brain volumes for each of the three runs were collected per participant. Field-maps were acquired for field unwarping.

**MRI pre-processing**

Functional data was first pre-processed using FSL-FEAT (version 6.00). Pre-processing included motion correction using MCFLIRT^45^, brain extraction using BET^46^, temporal high-pass filtering with a cut-off of 119 s and spatial smoothing using a Gaussian kernel with a FWHM of 3 mm. Field maps were used for distortion correction of functional data.

For each participant, we calculated a midspace between the three functional runs, i.e., the average space in which the images are minimally reorientated. Each functional run was then aligned to the midspace and registered to structural images (within-subject) using FMRIB’s Linear Image Registration Tool (FLIRT), and optimised using Boundary-Based Registration^47^. Where specified, functional and structural data were transformed to MNI152 space using FMRIB’s Nonlinear Registration Tool (FNRIT^48^).

**Group-level visualisations**

Prior to group-level visualisations, participant information regarding hand dominance (controls) and deprived hemisphere (one-handed participants) were used to sagittal-flip raw pre-processed data, such that the brain activity corresponding to the non-dominant/missing hand is always represented in the left hemisphere (note that a similar proportion of participants were flipped across groups). Group-level statistical parameter maps were then created with a threshold-free cluster enhancement (TFCE) approach using FSL’s Randomise tool^58^. TFCE is a nonparametric, permutation-based method for cluster formation, and has been shown to demonstrate improved sensitivity when compared to typical thresholding methods^59^. Group activity mixed-effect maps were calculated for each fixed-effect (i.e., averaged across the three functional runs) parameter estimate of a face movement (forehead, lips, tongue) contrasted to baseline. Prior to permutation (*n* = 5000), parameter estimates were masked with a sensorimotor mask, defined as the precentral and postcentral gyrus from the Harvard Cortical Atlas. A family-wise error correction of *p* < 0.05 and variance smoothing of 5 mm (as recommended for datasets with less than 20 participants) were used. Resulting clusters were thresholded at *p* < .001 and projected to a group cortical surface^60^ using Connectome Workbench (v1.4.2), and activity is visualised in Brodmann Areas 1, 2, 3a, 3b and 4.

As well as activity maps, we also visualised the winner-takes-all output at the group-level. Here the ‘winners’ for each face movement within the S1 ROI (the hand and face region combined) in MNI152 space were concatenated into a single volume per group to produce a consistency map for the individual movements (i.e., how many participants maximally activated the same voxel when moving a given facial part). Resulting consistency maps were then projected to a group cortical surface^61^ using Connectome Workbench (v1.4.2) for visualisation only.

**Validation**

***Procedure***

Two two-handed individuals took part in this validation procedure (aged 33 and 29, 2 women, 1 left-handed). These individuals underwent two sessions of the sensorimotor active task used in the main analyses (hereafter Active1 and Active2), and allowed us to assess the consistency of our data.

***Functional MRI data acquisition and analysis***

Functional and anatomical MRI data for the Active2 session was obtained with the same MRI parameters as for the Active1 session (collected on the same scanner as the data collected for the main analyses), but on a different 3T Prisma scanner. Functional data was pre-processed and analysed as described for the main analyses.

***Results***

A significant positive relationship between the two active sessions was found for both participants (see Supplementary Figure 1B), indicating a relatively stable representational pattern of facial activity in the face region across time (see Supplementary Figure 1A). This indicates a broadly similar representational structure of the face between the two sessions, highlight the stability of the facial representation across time.

***Supplementary Figue 1. Comparison of active1 versus active2 stimulation of the face for two control participants in the non-dominant hemisphere. (A)*** *Representational Dissmialarity Matrices (RDMs) for two control participants in the face region of interest (see Methods; Multivariate representational analysis) who completed two active motor paradigm (see Methods; Functional MRI sensorimotor task) sessions ~12 months apart (see Supplementary Methods; Validation). Greater dissimilarity between activity patterns for the pairwise comparison indicates an increased ability to discriminate between the two facial movements/stimulations in the face region, i.e., there is a greater amount of facial information content. Smaller dissimilarity values indicated a reduced ability to discriminate between the two face parts.* ***(B)*** *Pearson’s correlations examining the relationship between face-face and face-thumb dissimilarity values for both the first and second active motor paradigm sessions.*

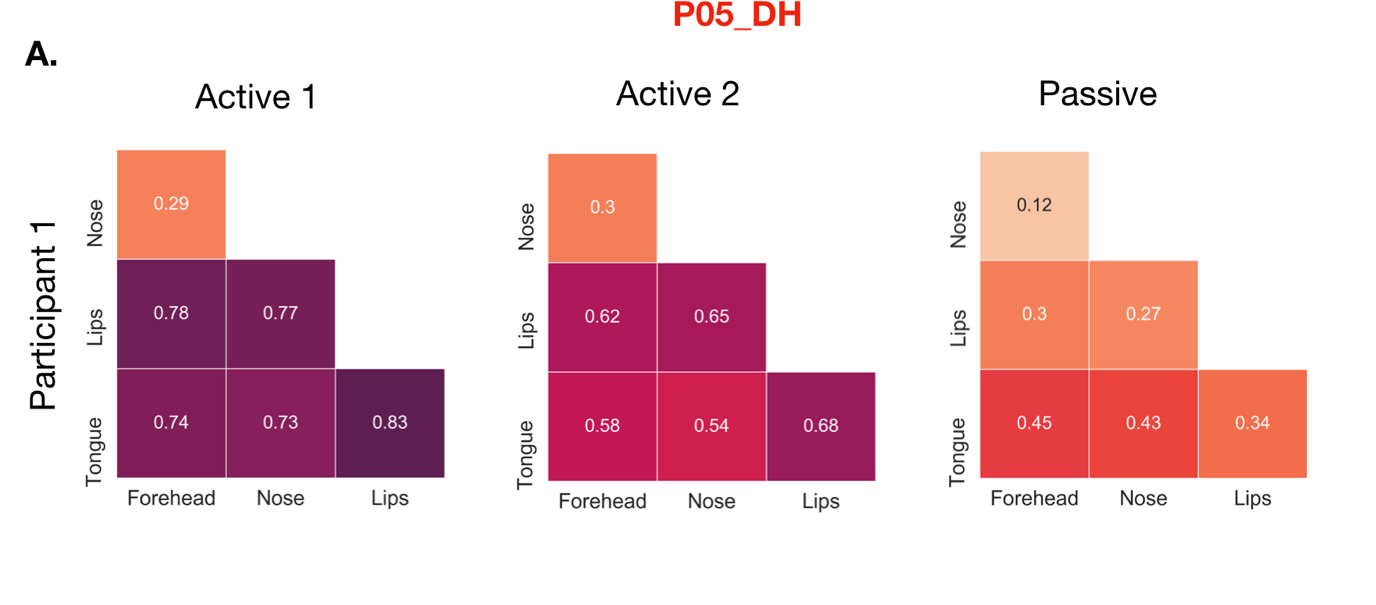

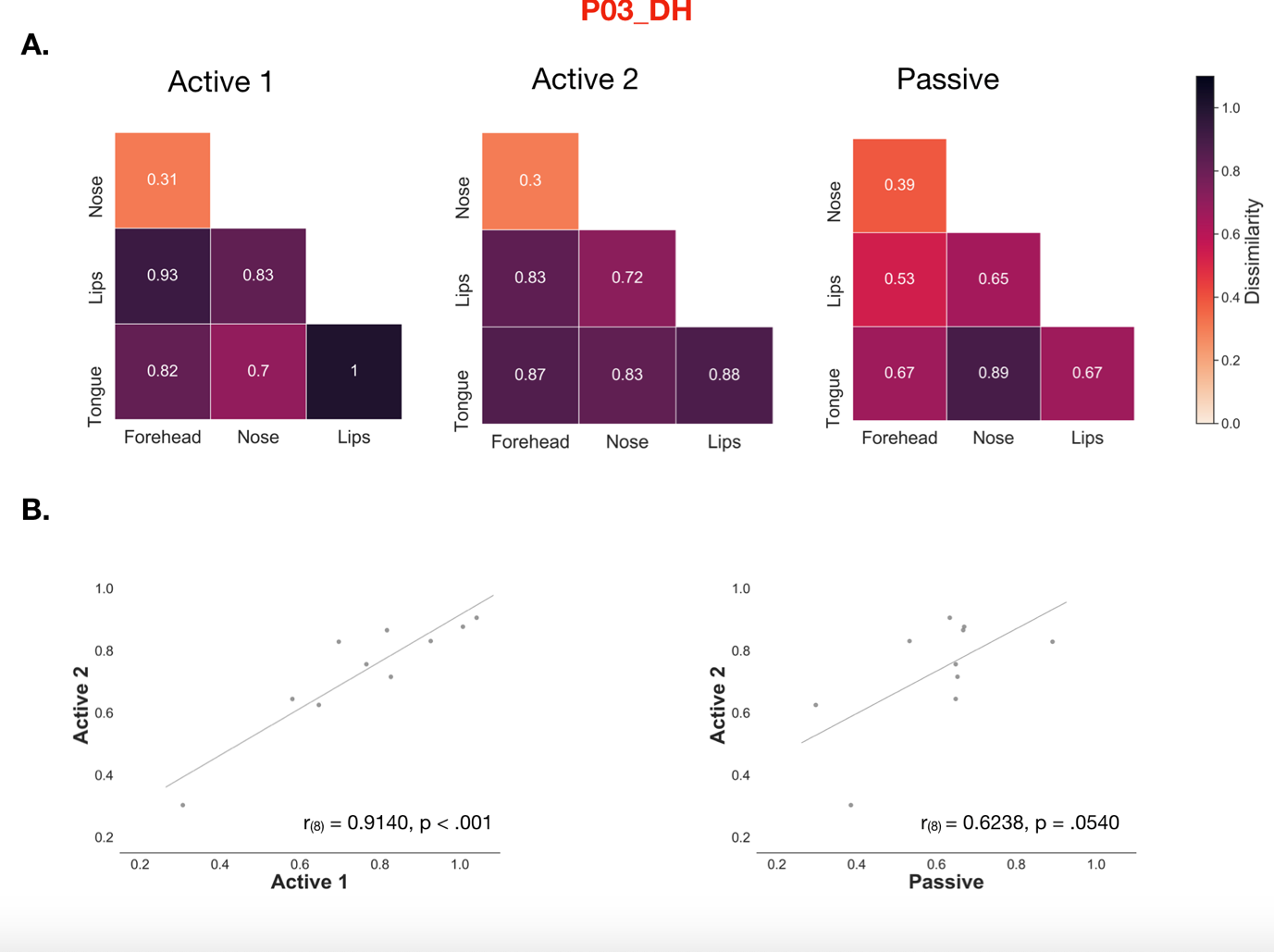

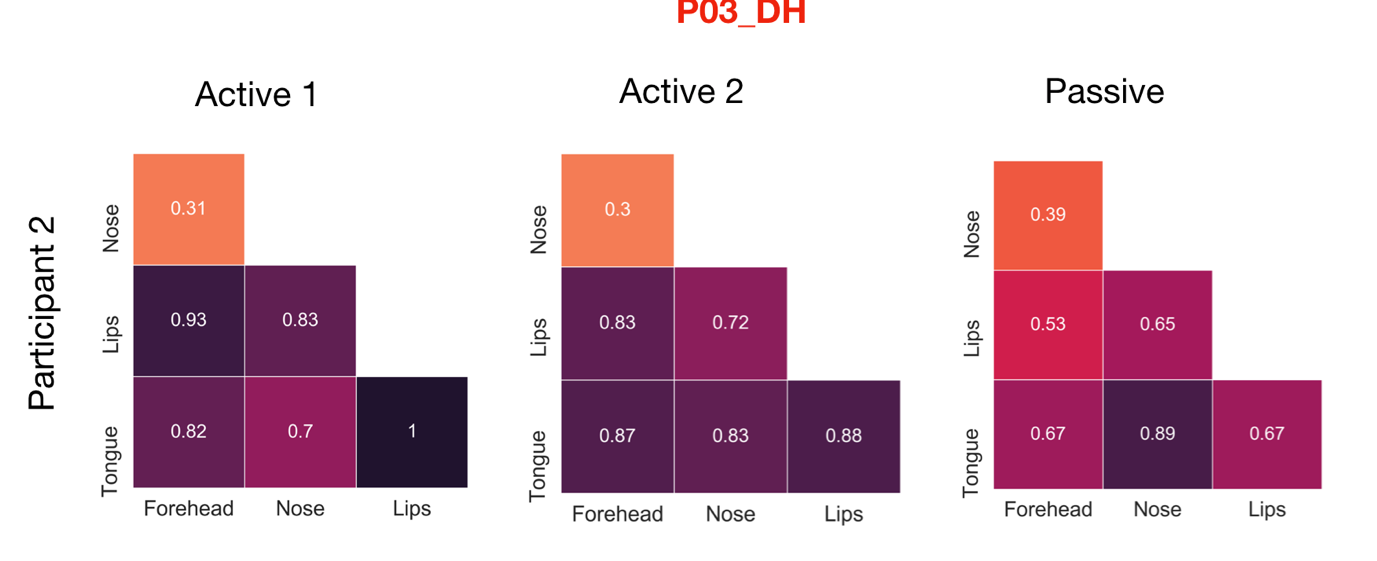

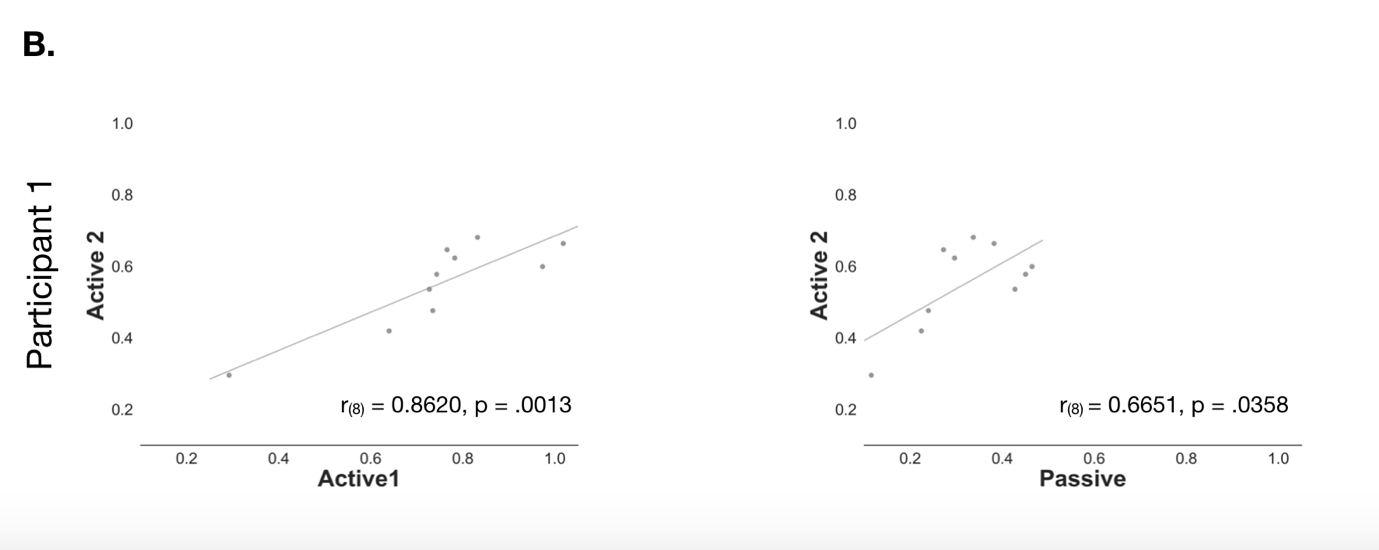

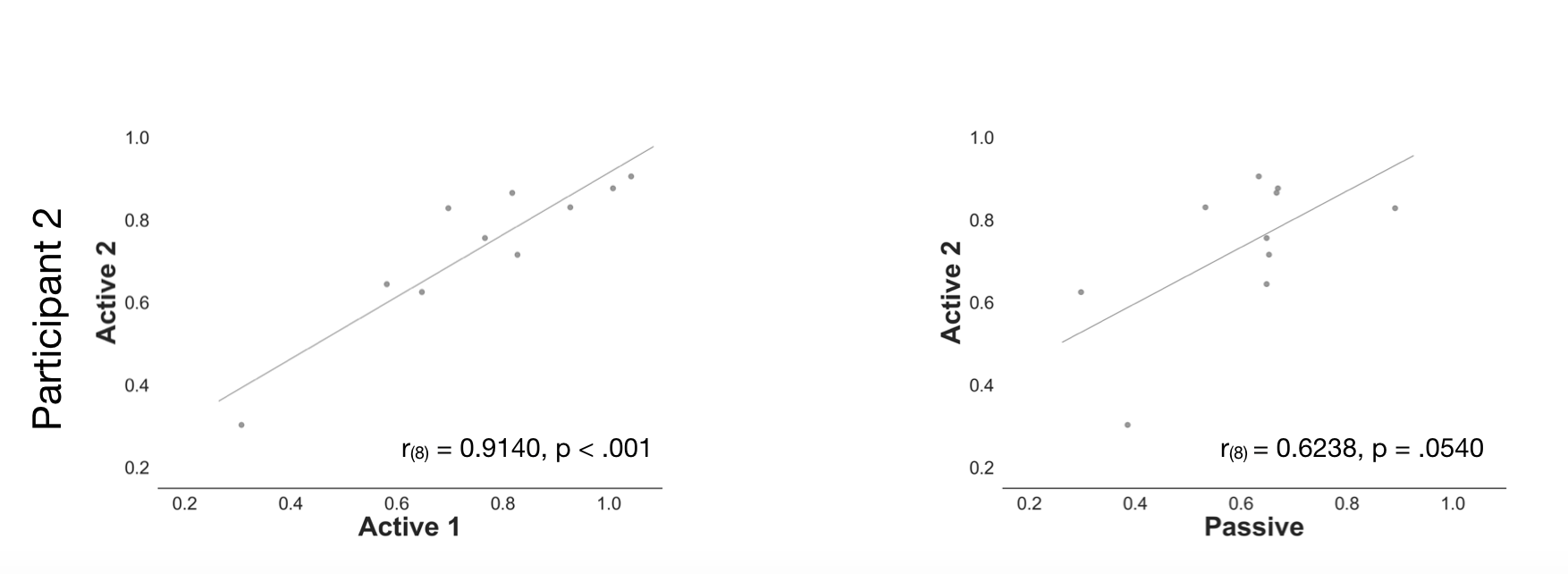

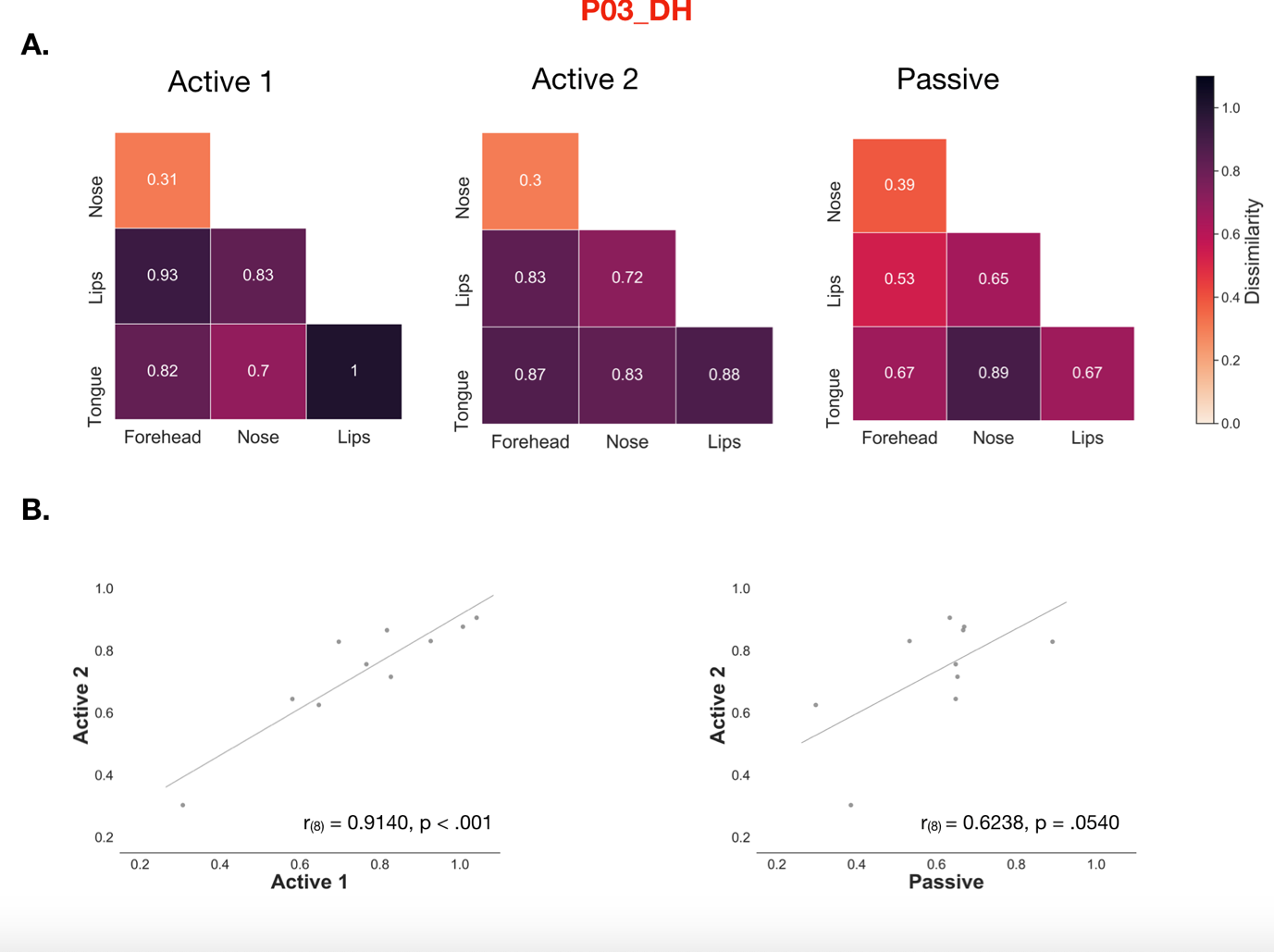

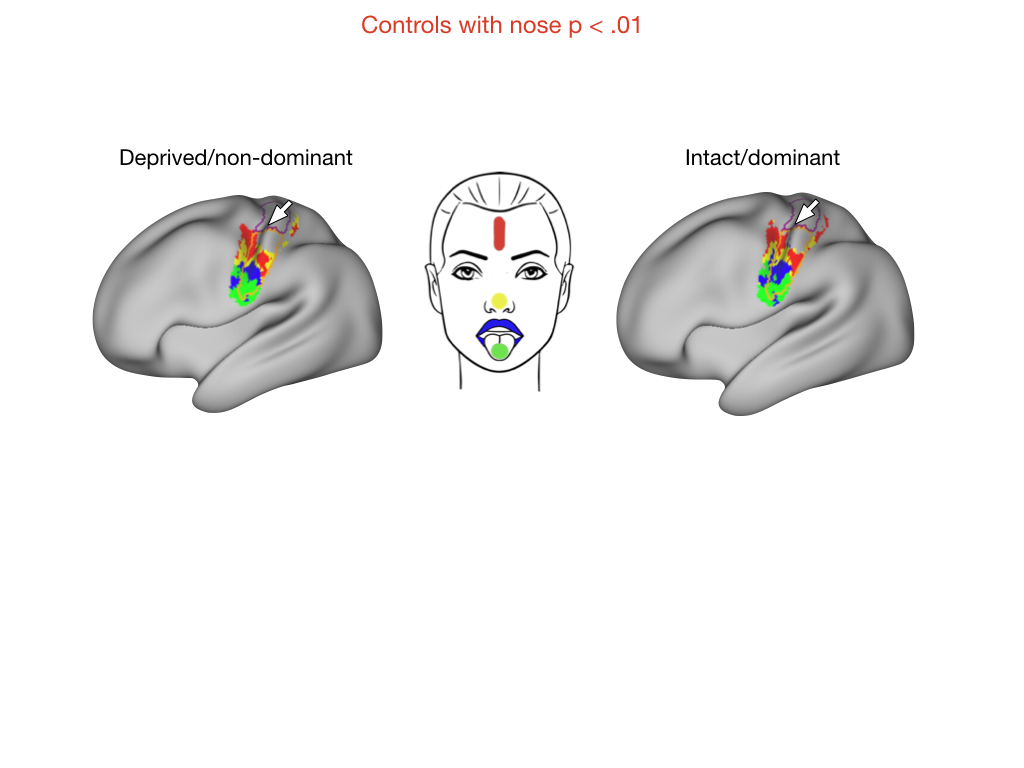

***Supplementary Figure 2. Group-level activity maps of control participants with nose movement.*** *Group average activity for the forehead (red), nose (yellow), lips (blue) and tongue (green) movements, contrasted to rest, in the deprived/non-dominant hemisphere and intact/dominant hemisphere for controls (n = 22). All clusters were created using a threshold-free cluster enhancement procedure with a sensorimotor pre-threshold mask (defined using the Harvard Cortical Atlas), and thresholded at p < .01. The hand and face ROIs are outlined in purple and orange respectively, and the central sulcus is denoted with a white arrow.*

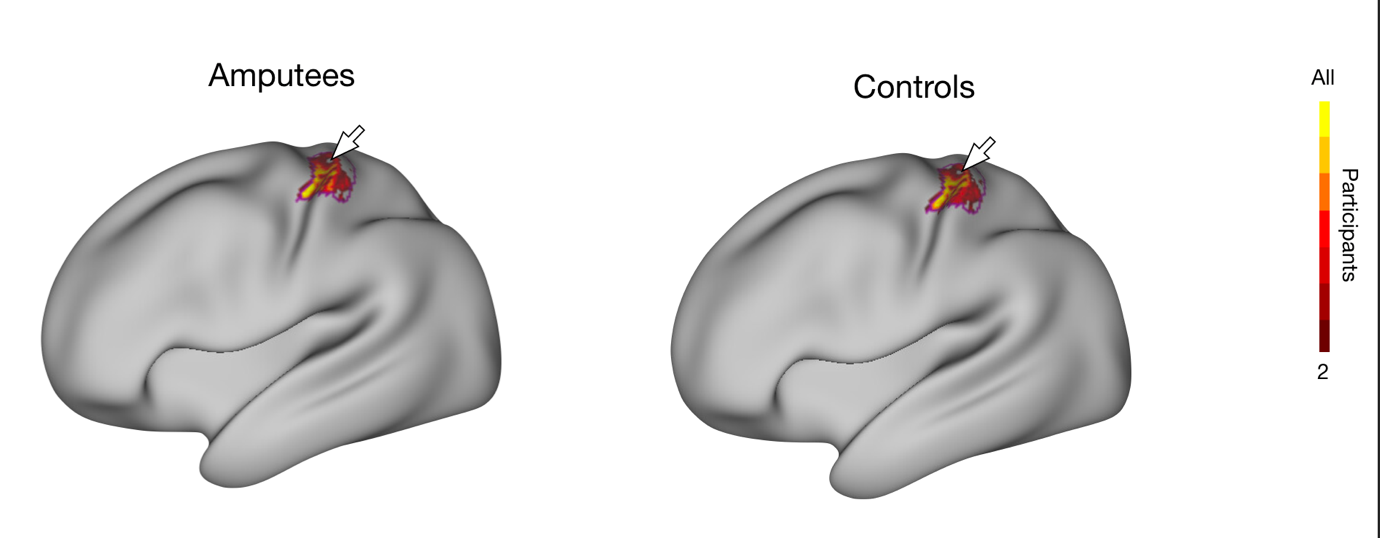

***Supplementary Figure 3. Phantom and non-dominant thumb representation in the deprived hemisphere of amputees and controls.*** *Group-level consistency map for the phantom/non-dominant thumb in the hand ROI for amputees (n = 17) and controls (n = 22). Note the percentage of surface area coverage of the deprived/non-dominant hand ROI was not significantly different for the phantom (M=69.448%; SE=3.752%) compared to the non-dominant thumb of controls (M=63.989%; SE=3.594 %; U=225.000, p=0.292, d=0.203, BF_10_=0.579). The colour gradient represents participant agreement for maximally activating that particular voxel, relative to the face movements (winner-takes-all approach). The hand ROI is outlined in purple and central sulcus denoted by the white arrow.*

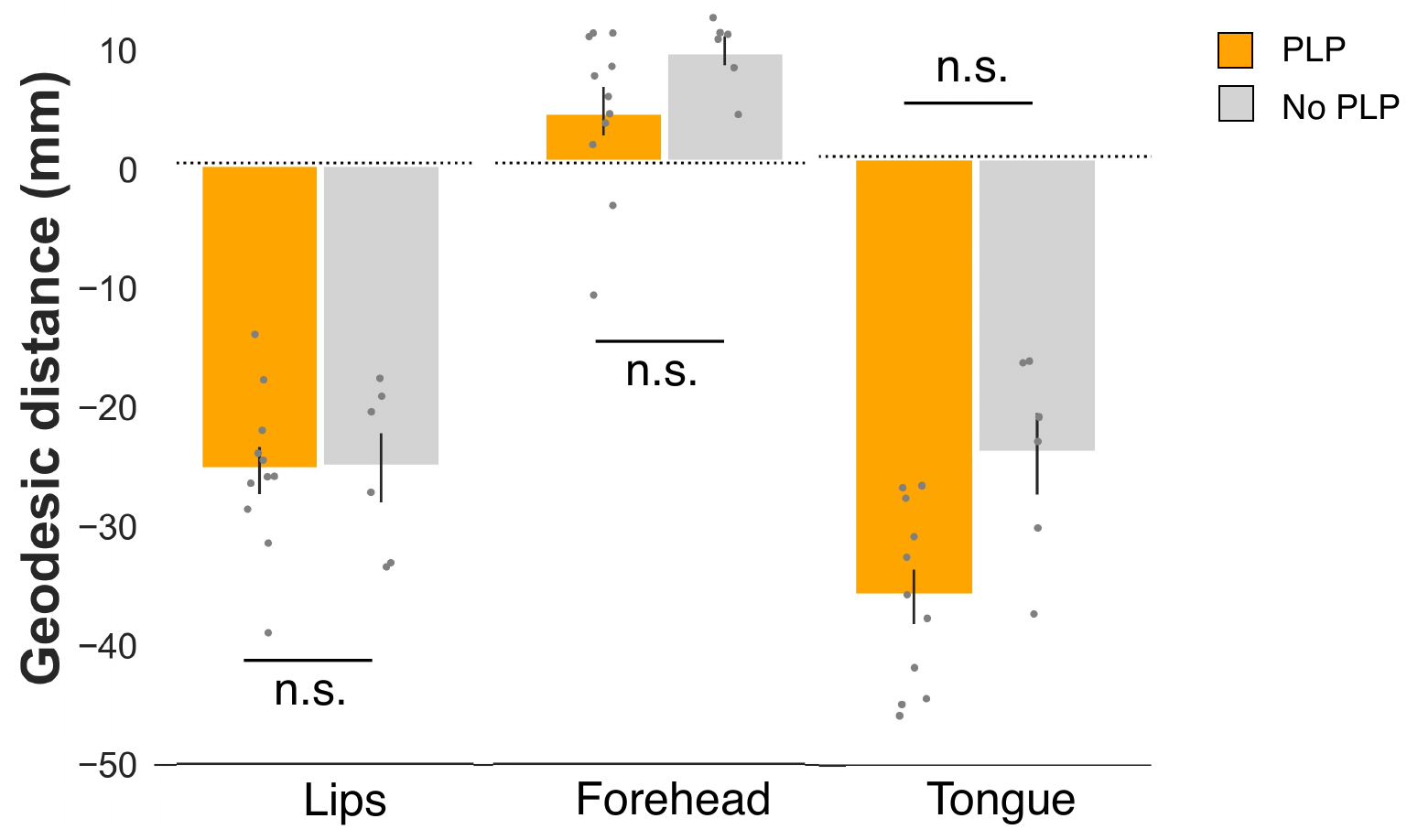

***Supplementary Figure 4. Comparison of cortical distances in the deprived hemisphere of amputees.*** *Cortical (geodesic) distances were compared between amputees who reported the presence of PLP (n=11; orange) and amputees without PLP (n=6; grey) using a one-tailed Mann-Whitney test. Non-significant differences were found for the lips (t_(15)_=-0.068, p=0.527, d=-0.035, BF_10_=0.414), forehead (t_(15)_=-1.720, p=0.947, d=-0.873, BF_10_=0.203) and tongue (t_(15)_=-3.018, p=0.996, d=-1.532, BF_10_=0.156). Contrary to popular theories of brain plasticity and phantom limb pain (PLP; see Introduction), these results demonstrate that individuals with PLP do not exhibit greater instances of cortical remapping in the deprived hemisphere of the tested facial parts (including both the traditional marker of plasticity – the lips – and the cortical neighbour – the forehead).*

***Supplementary Table 1. Statistics for comparisons between amputees and controls***

***(A) Main effects and interaction for comparison of geodesic distances between amputees and controls for the lips.***

| **Within Subjects Effects** | | | | | | | | | | | | |
| --- | --- | --- | --- | --- | --- | --- | --- | --- | --- | --- | --- | --- |
| **Cases** | | **Sum of Squares** | | **df** | | **Mean Square** | | **F** | | **p** | | **η²** |
| Hemisphere |  | 0.003 |  | 1 |  | 0.003 |  | 1.910e -4 |  | 0.989 |  | 1.338e -6 |
| Hemisphere ✻ group |  | 0.053 |  | 1 |  | 0.053 |  | 0.003 |  | 0.954 |  | 2.413e -5 |
| Hemisphere ✻ brainVol |  | 0.092 |  | 1 |  | 0.092 |  | 0.006 |  | 0.939 |  | 4.189e -5 |
| Residuals |  | 553.654 |  | 36 |  | 15.379 |  |  |  |  |  |  |
| *Note.*  Type III Sum of Squares | | | | | | | | | | | | |

| **Between Subjects Effects** | | | | | | | | | | | | |
| --- | --- | --- | --- | --- | --- | --- | --- | --- | --- | --- | --- | --- |
| **Cases** | | **Sum of Squares** | | **df** | | **Mean Square** | | **F** | | **p** | | **η²** |
| group |  | 67.225 |  | 1 |  | 67.225 |  | 1.821 |  | 0.186 |  | 0.031 |
| brainVol |  | 245.659 |  | 1 |  | 245.659 |  | 6.655 |  | 0.014 |  | 0.112 |
| Residuals |  | 1328.857 |  | 36 |  | 36.913 |  |  |  |  |  |  |
| *Note.*  Type III Sum of Squares | | | | | | | | | | | | |

***(B) Main effects and interaction for comparison of geodesic distances between amputees and controls for the forehead.***

| **Within Subjects Effects** | | | | | | | | | | | | |
| --- | --- | --- | --- | --- | --- | --- | --- | --- | --- | --- | --- | --- |
| **Cases** | | **Sum of Squares** | | **df** | | **Mean Square** | | **F** | | **p** | | **η²** |
| Hemisphere |  | 4.254 |  | 1 |  | 4.254 |  | 0.229 |  | 0.635 |  | 0.003 |
| Hemisphere ✻ group |  | 17.380 |  | 1 |  | 17.380 |  | 0.935 |  | 0.340 |  | 0.011 |
| Hemisphere ✻ brainVol |  | 2.752 |  | 1 |  | 2.752 |  | 0.148 |  | 0.703 |  | 0.002 |
| Residuals |  | 668.911 |  | 36 |  | 18.581 |  |  |  |  |  |  |
| Note.  Type III Sum of Squares | | | | | | | | | | | | |

| **Between Subjects Effects** | | | | | | | | | | | | |
| --- | --- | --- | --- | --- | --- | --- | --- | --- | --- | --- | --- | --- |
| **Cases** | | **Sum of Squares** | | **df** | | **Mean Square** | | **F** | | **p** | | **η²** |
| group |  | 14.117 |  | 1 |  | 14.117 |  | 0.634 |  | 0.431 |  | 0.009 |
| brainVol |  | 92.715 |  | 1 |  | 92.715 |  | 4.164 |  | 0.049 |  | 0.058 |
| Residuals |  | 801.643 |  | 36 |  | 22.268 |  |  |  |  |  |  |
| Note.  Type III Sum of Squares | | | | | | | | | | | | |

***(C) Main effects and interaction for comparison of geodesic distances between amputees and controls for the tongue.***

| **Within Subjects Effects** | | | | | | | | | | | | |
| --- | --- | --- | --- | --- | --- | --- | --- | --- | --- | --- | --- | --- |
| **Cases** | | **Sum of Squares** | | **df** | | **Mean Square** | | **F** | | **p** | | **η²** |
| Hemisphere |  | 0.164 |  | 1 |  | 0.164 |  | 0.011 |  | 0.918 |  | 2.843e -5 |
| Hemisphere ✻ group |  | 38.760 |  | 1 |  | 38.760 |  | 2.546 |  | 0.119 |  | 0.007 |
| Hemisphere ✻ brainVol |  | 0.794 |  | 1 |  | 0.794 |  | 0.052 |  | 0.821 |  | 1.374e -4 |
| Residuals |  | 548.070 |  | 36 |  | 15.224 |  |  |  |  |  |  |
| Note.  Type III Sum of Squares | | | | | | | | | | | | |

| **Between Subjects Effects** | | | | | | | | | | | | |
| --- | --- | --- | --- | --- | --- | --- | --- | --- | --- | --- | --- | --- |
| **Cases** | | **Sum of Squares** | | **df** | | **Mean Square** | | **F** | | **p** | | **η²** |
| group |  | 106.146 |  | 1 |  | 106.146 |  | 0.797 |  | 0.378 |  | 0.018 |
| brainVol |  | 285.276 |  | 1 |  | 285.276 |  | 2.142 |  | 0.152 |  | 0.049 |
| Residuals |  | 4795.420 |  | 36 |  | 133.206 |  |  |  |  |  |  |
| Note.  Type III Sum of Squares | | | | | | | | | | | | |

***Supplementary Table 2. Statistics for comparisons between one-handers and controls***

***(A) Main effects and interaction for comparison of geodesic distances of the lips between one-handers and controls.***

| **Within Subjects Effects** | | | | | | | | | | | | |
| --- | --- | --- | --- | --- | --- | --- | --- | --- | --- | --- | --- | --- |
| **Cases** | | **Sum of Squares** | | **df** | | **Mean Square** | | **F** | | **p** | | **η²** |
| Hemisphere |  | 4.987 |  | 1 |  | 4.987 |  | 0.290 |  | 0.593 |  | 0.002 |
| Hemisphere ✻ group |  | 93.236 |  | 1 |  | 93.236 |  | 5.419 |  | 0.025 |  | 0.032 |
| Hemisphere ✻ brainVol |  | 0.920 |  | 1 |  | 0.920 |  | 0.053 |  | 0.818 |  | 3.120e -4 |
| Residuals |  | 688.253 |  | 40 |  | 17.206 |  |  |  |  |  |  |
| *Note.*  Type III Sum of Squares | | | | | | | | | | | | |

| **Between Subjects Effects** | | | | | | | | | | | | |
| --- | --- | --- | --- | --- | --- | --- | --- | --- | --- | --- | --- | --- |
| **Cases** | | **Sum of Squares** | | **df** | | **Mean Square** | | **F** | | **p** | | **η²** |
| group |  | 7.777 |  | 1 |  | 7.777 |  | 0.194 |  | 0.662 |  | 0.003 |
| brainVol |  | 550.077 |  | 1 |  | 550.077 |  | 13.725 |  | < .001 |  | 0.187 |
| Residuals |  | 1603.114 |  | 40 |  | 40.078 |  |  |  |  |  |  |
| *Note.*  Type III Sum of Squares | | | | | | | | | | | | |

***(B) Main effects and interaction for comparison of geodesic distances between one-handers and controls for the forehead.***

| **Within Subjects Effects** | | | | | | | | | | | | |
| --- | --- | --- | --- | --- | --- | --- | --- | --- | --- | --- | --- | --- |
| **Cases** | | **Sum of Squares** | | **df** | | **Mean Square** | | **F** | | **p** | | **η²** |
| Hemisphere |  | 0.070 |  | 1 |  | 0.070 |  | 0.005 |  | 0.941 |  | 3.886e -5 |
| Hemisphere ✻ group |  | 106.153 |  | 1 |  | 106.153 |  | 8.287 |  | 0.006 |  | 0.059 |
| Hemisphere ✻ brainVol |  | 0.134 |  | 1 |  | 0.134 |  | 0.010 |  | 0.919 |  | 7.449e -5 |
| Residuals |  | 512.392 |  | 40 |  | 12.810 |  |  |  |  |  |  |
| Note.  Type III Sum of Squares | | | | | | | | | | | | |

| **Between Subjects Effects** | | | | | | | | | | | | |
| --- | --- | --- | --- | --- | --- | --- | --- | --- | --- | --- | --- | --- |
| **Cases** | | **Sum of Squares** | | **df** | | **Mean Square** | | **F** | | **p** | | **η²** |
| group |  | 68.526 |  | 1 |  | 68.526 |  | 2.575 |  | 0.116 |  | 0.038 |
| brainVol |  | 50.946 |  | 1 |  | 50.946 |  | 1.914 |  | 0.174 |  | 0.028 |
| Residuals |  | 1064.519 |  | 40 |  | 26.613 |  |  |  |  |  |  |
| Note.  Type III Sum of Squares | | | | | | | | | | | | |

***(C) Main effects and interaction for comparison of geodesic distances between one-handers and controls for the tongue.***

| **Within Subjects Effects** | | | | | | | | | | | | |
| --- | --- | --- | --- | --- | --- | --- | --- | --- | --- | --- | --- | --- |
| **Cases** | | **Sum of Squares** | | **df** | | **Mean Square** | | **F** | | **p** | | **η²** |
| Hemisphere |  | 39.972 |  | 1 |  | 39.972 |  | 1.493 |  | 0.229 |  | 0.007 |
| Hemisphere ✻ group |  | 191.346 |  | 1 |  | 191.346 |  | 7.149 |  | 0.011 |  | 0.032 |
| Hemisphere ✻ brainVol |  | 25.683 |  | 1 |  | 25.683 |  | 0.960 |  | 0.333 |  | 0.004 |
| Residuals |  | 1070.656 |  | 40 |  | 26.766 |  |  |  |  |  |  |
| Note.  Type III Sum of Squares | | | | | | | | | | | | |

| **Between Subjects Effects** | | | | | | | | | | | | |
| --- | --- | --- | --- | --- | --- | --- | --- | --- | --- | --- | --- | --- |
| **Cases** | | **Sum of Squares** | | **df** | | **Mean Square** | | **F** | | **p** | | **η²** |
| group |  | 174.848 |  | 1 |  | 174.848 |  | 1.732 |  | 0.196 |  | 0.029 |
| brainVol |  | 441.349 |  | 1 |  | 441.349 |  | 4.373 |  | 0.043 |  | 0.074 |
| Residuals |  | 4037.482 |  | 40 |  | 100.937 |  |  |  |  |  |  |
| Note.  Type III Sum of Squares | | | | | | | | | | | | |

***Supplementary Table 3. Results from the linear mixed model used to explore differences in face-thumb pairwise distances in the hand ROI for amputees and controls.***

| Fixed Effect Omnibus tests | | | | | | | | |
| --- | --- | --- | --- | --- | --- | --- | --- | --- |
|  | | **F** | | **Num df** | | **Den df** | | **p** |
| Group |  | 2.2211 |  | 1 |  | 37.0 |  | 0.145 |
| Hemisphere |  | 1.8462 |  | 1 |  | 259.0 |  | 0.175 |
| faceThumbPairs |  | 3.4735 |  | 3 |  | 259.0 |  | 0.017 |
| Group ✻ Hemisphere |  | 0.0571 |  | 1 |  | 259.0 |  | 0.811 |
| Group ✻ faceThumbPairs |  | 0.7404 |  | 3 |  | 259.0 |  | 0.529 |
| Hemisphere ✻ faceThumbPairs |  | 0.3653 |  | 3 |  | 259.0 |  | 0.778 |
| Group ✻ Hemisphere ✻ faceThumbPairs |  | 0.5692 |  | 3 |  | 259.0 |  | 0.636 |
| Note. Satterthwaite method for degrees of freedom | | | | | | | | |

***Supplementary Table 4. Results from the linear mixed model used to explore differences in face-face pairwise distances in the hand ROI for amputees, one-handers and controls.***

| Fixed Effect Omnibus tests | | | | | | | | |
| --- | --- | --- | --- | --- | --- | --- | --- | --- |
|  | | **F** | | **Num df** | | **Den df** | | **p** |
| Group |  | 2.409 |  | 2 |  | 56.0 |  | 0.099 |
| Hemisphere |  | 27.059 |  | 1 |  | 627.0 |  | < .001 |
| FacePairs |  | 22.380 |  | 5 |  | 627.0 |  | < .001 |
| Age |  | 0.286 |  | 1 |  | 56.0 |  | 0.595 |
| Group ✻ Hemisphere |  | 14.544 |  | 2 |  | 627.0 |  | < .001 |
| Group ✻ FacePairs |  | 0.881 |  | 10 |  | 627.0 |  | 0.551 |
| Hemisphere ✻ FacePairs |  | 1.056 |  | 5 |  | 627.0 |  | 0.384 |
| Group ✻ Hemisphere ✻ FacePairs |  | 0.572 |  | 10 |  | 627.0 |  | 0.837 |
| Note. Satterthwaite method for degrees of freedom | | | | | | | | |

***Supplementary Table 5. Results from the linear mixed model used to explore differences in face-face pairwise distances in the face ROI for amputees, one-handers and controls.***

| Fixed Effect Omnibus tests | | | | | | | | |
| --- | --- | --- | --- | --- | --- | --- | --- | --- |
|  | | **F** | | **Num df** | | **Den df** | | **p** |
| Group |  | 1.010 |  | 2 |  | 56.0 |  | 0.371 |
| Hemisphere |  | 1.301 |  | 1 |  | 627.0 |  | 0.254 |
| FacePairs |  | 268.322 |  | 5 |  | 627.0 |  | < .001 |
| Age |  | 0.282 |  | 1 |  | 56.0 |  | 0.598 |
| Group ✻ Hemisphere |  | 0.626 |  | 2 |  | 627.0 |  | 0.535 |
| Group ✻ FacePairs |  | 1.462 |  | 10 |  | 627.0 |  | 0.150 |
| Hemisphere ✻ FacePairs |  | 0.222 |  | 5 |  | 627.0 |  | 0.953 |
| Group ✻ Hemisphere ✻ FacePairs |  | 0.136 |  | 10 |  | 627.0 |  | 0.999 |
| Note. Satterthwaite method for degrees of freedom | | | | | | | | |
